# Transcriptional regulation of amino acid metabolism by KDM2B, in the context of ncPRC1.1 and in concert with MYC and ATF4

**DOI:** 10.1101/2023.07.07.548031

**Authors:** Evangelia Chavdoula, Vollter Anastas, Alessandro La Ferlita, Julian Aldana, Giuseppe Carota, Mariarita Spampinato, Burak Soysal, Ilaria Cosentini, Sameer Parashar, Anuvrat Sircar, Giovanni Nigita, Lalit Sehgal, Michael A. Freitas, Philip N. Tsichlis

## Abstract

**Introduction:** KDM2B encodes a JmjC domain-containing histone lysine demethylase, which functions as an oncogene in several types of tumors, including TNBC. This study was initiated to address the cancer relevance of the results of our earlier work, which had shown that overexpression of KDM2B renders mouse embryonic fibroblasts (MEFs) resistant to oxidative stress by regulating antioxidant mechanisms.

**Methods:** We mainly employed a multi-omics strategy consisting of RNA-Seq, quantitative TMT proteomics, Mass-spectrometry-based global metabolomics, ATAC-Seq and ChIP-seq, to explore the role of KDM2B in the resistance to oxidative stress and intermediary metabolism. These data and data from existing patient datasets were analyzed using bioinformatic tools, including exon-intron-split analysis (EISA), FLUFF and clustering analyses. The main genetic strategy we employed was gene silencing with shRNAs. ROS were measured by flow cytometry, following staining with CellROX and various metabolites were measured with biochemical assays, using commercially available kits. Gene expression was monitored with qRT-PCR and immunoblotting, as indicated.

**Results:** The knockdown of KDM2B in basal-like breast cancer cell lines lowers the levels of GSH and sensitizes the cells to ROS inducers, GSH targeting molecules, and DUB inhibitors. To address the mechanism of GSH regulation, we knocked down KDM2B in MDA-MB-231 cells and we examined the effects of the knockdown, using a multi-omics strategy. The results showed that KDM2B, functioning in the context of ncPRC1.1, regulates a network of epigenetic and transcription factors, which control a host of metabolic enzymes, including those involved in the SGOC, glutamate, and GSH metabolism. They also showed that KDM2B enhances the chromatin accessibility and expression of MYC and ATF4, and that it binds in concert with MYC and ATF4, the promoters of a large number of transcriptionally active genes, including many, encoding metabolic enzymes. Additionally, MYC and ATF4 binding sites were enriched in genes whose accessibility depends on KDM2B, and analysis of a cohort of TNBCs expressing high or low levels of KDM2B, but similar levels of MYC and ATF4 identified a subset of MYC targets, whose expression correlates with the expression of KDM2B. Further analyses of basal-like TNBCs in the same cohort, revealed that tumors expressing high levels of all three regulators exhibit a distinct metabolic signature that carries a poor prognosis.

**Conclusions:** The present study links KDM2B, ATF4, and MYC in a transcriptional network that regulates the expression of multiple metabolic enzymes, including those that control the interconnected SGOC, glutamate, and GSH metabolic pathways. The co-occupancy of the promoters of many transcriptionally active genes, by all three factors, the enrichment of MYC binding sites in genes whose chromatin accessibility depends on KDM2B, and the correlation of the levels of KDM2B with the expression of a subset of MYC target genes in tumors that express similar levels of MYC, suggest that KDM2B regulates both the expression and the transcriptional activity of MYC. Importantly, the concerted expression of all three factors also defines a distinct metabolic subset of TNBCs with poor prognosis. Overall, this study identifies novel mechanisms of SGOC regulation, suggests novel KDM2B-dependent metabolic vulnerabilities in TNBC, and provides new insights into the role of KDM2B in the epigenetic regulation of transcription.

**Graphical Abstract:** 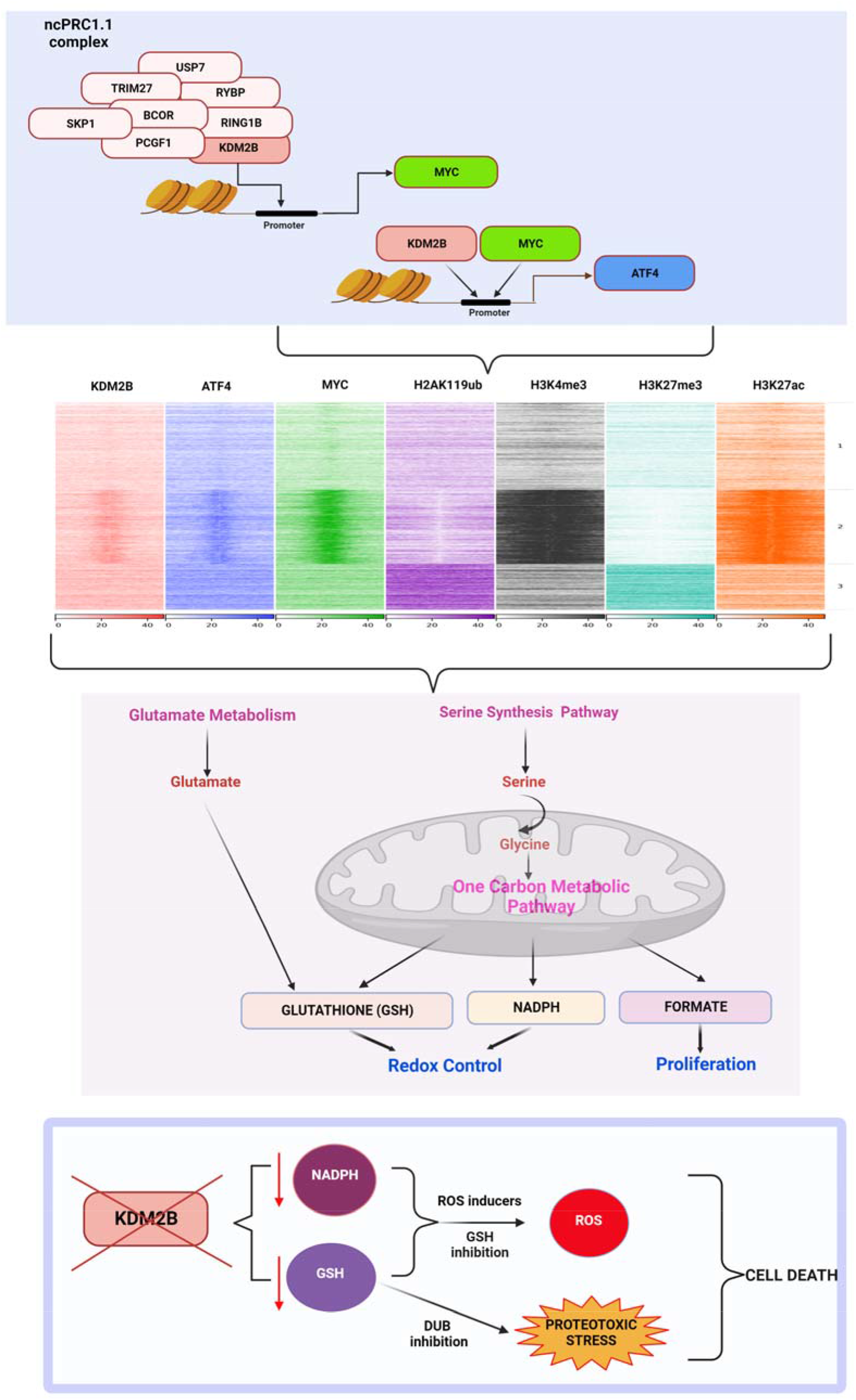

**Highlights:** The knockdown of KDM2B in basal-like breast cancer cell lines lowers the levels of GSH and sensitizes the cells to ROS inducers, GSH targeting molecules, and DUB inhibitors.

KDM2B regulates intermediary metabolism by targeting the expression of a host of metabolic enzymes, including those in the SGOC, glutamate, and GSH metabolism.

KDM2B enhances chromatin accessibility of MYC and ATF4 and promotes their expression.

MYC and ATF4 binding sites are enriched in the promoters of genes whose accessibility depends on KDM2B.

KDM2B functioning in the context of ncPRC1.1, binds the promoters of transcriptionally active genes, including those encoding KDM2B-regulated metabolic enzymes, in concert with MYC and ATF4.

Basal-like TNBCs expressing high levels of all three regulators, exhibit a distinct metabolic signature that is associated with poor prognosis.

## Introduction

*KDM2B* (also known as *NDY1, FBXL10, JHDM1B* or *Fbl10*), encodes a JmjC domain-containing histone lysine demethylase, which targets histone H3K36me2/me1 and perhaps histone H3K4me3 [1, 2, 3, 4] and histone H3K79 me3/me2 [5]. KDM2B functions as an oncogene in several types of tumors. Its role in oncogenesis was originally observed in Moloney MuLV-induced rodent T cell lymphomas, where it is activated by provirus integration in the vicinity of its TSS [2]. Subsequently, its oncogenic role was extended to human lymphoid and myeloid malignancies [6, 7], and to bladder [8] and pancreatic cancer [9], basal-like breast cancer [10], gliomas [11] and prostate cancer [12]. Its oncogenic potential is mediated by multiple mechanisms, including stimulation of cell proliferation [7, 9, 10, 13, 14], inhibition of cellular senescence [2,4] inhibition of apoptosis [8,10], resistance to oxidative stress [15], regulation of cellular metabolism [16], regulation of stem cell reprogramming [17,18], differentiation [6], and cancer stem cell maintenance [10]. In addition to the inhibition of cellular senescence, which was the first observation providing mechanistic insights into the role of KDM2B in oncogenesis [2,4], our studies addressing the effects of the KDM2B knockdown in a set of ten cancer cell lines, showed that KDM2B promotes cell cycle progression in all and inhibits apoptosis in some [10].

The focus of this report is on the role of KDM2B in the metabolism of basal-like TNBC, which comprises 15-20% of all breast cancers and is more prevalent in younger women (age <40 years). These tumors tend to express high levels of KDM2B, they do not express estrogen and progesterone receptors, and they are negative for the amplification of epidermal growth factor receptor 2 [19]. Most basal-like breast cancers exhibit an aggressive clinical course with poor overall survival and early relapse after treatment [19, 20]. The poor prognosis of these tumors is partially due to the lack of effective targeted therapies [20] and the goal of this work is to improve our understanding of the biology of the disease, which may lead to novel and more effective therapeutic strategies. Our earlier studies addressing the role of KDM2B in metabolism revealed that it upregulates a set of antioxidant genes, including Aass, Nqo1, Prdx4 and Serpinb1b, and protects cells from oxidative stress [15]. More recent studies showed that KDM2B downregulates PDH and shifts metabolism toward aerobic glycolysis and glutaminolysis. It upregulates ATCase (or CAD), and promotes pyrimidine biosynthesis [16].

Multiomic strategies employed in the course of this study, revealed that KDM2B exerts global effects on cellular metabolism. One of its major targets is the One-Carbon (1C) metabolism pathway, which controls many processes, including the methylation of cellular metabolites and macromolecules, nucleotide biosynthesis, and antioxidant defenses, whose reprogramming is a common feature of oncogenesis [21, 22, 23, 24, 25]. The primary sources of 1C units are serine and glycine. Serine is produced primarily from the glycolytic intermediate 3PG via the Serine Synthesis Pathway (SSP). Glycine is produced primarily from serine, which donates a methyl group to THF, via a reversible reaction catalyzed by a pyridoxal phosphate-dependent enzyme. 1C metabolism couples the folate and methionine cycles of the SGOC pathway (SGOCP). The latter is compartmentalized and consists of two full pathways that operate in the mitochondria and the cytosol and are functionally linked via formate, which freely diffuses between the two compartments [23, 24, 25]. Although both branches of the pathway are active, cancer cells rely primarily on the mitochondrial branch [23, 26, 27, 28, 29]. SGOCP, in combination with the Pentose Phosphate Pathway (PPP), controls nucleotide biosynthesis, a process that is consistently activated in almost all tumors [23]. Several additional SGOCP-controlled metabolic modules are also activated in tumor subsets, rather than in all tumors, and their activation is context-dependent. One of them is redox maintenance, which depends on NADPH, the main reductant used in redox reactions, and GSH, the most abundant antioxidant, whose synthesis is under the control of the SGOC-linked transsulfuration pathway [24, 25, 30]. Importantly all modules of SGOCP are active in TNBC [29, 31, 32, 33, 34, 35, 36].

In addition to cancer, SGOCP has an important role in development, as evidenced by two observations: a) folate deficiency and genetic changes resulting in inhibition of the pathway give rise to mitochondrial dysfunction and spinal closure defects [37]; and b) SGOCP promotes stem cell self-renewal [38, 39]. Importantly, earlier studies had shown that KDM2B deficiency may also give rise to neural tube closure defects in mice [40] and that it promotes the self-renewal of normal and cancer stem cells [10, 17]. These observations, combined with the established oncogenic role of SGOCP, suggested a biologically important functional link between KDM2B and SGOCP and provided the rationale for studies exploring the role of KDM2B in SGOCP regulation.

The main transcriptional regulators of the genes encoding the enzymes involved in SGOCP are ATF4 [24, 25, 41, 42, 43] and the proto-oncogene c-MYC [22, 24, 25, 29, 44] and our data show that the expression of both, along with the expression of their enzyme-encoding gene targets is promoted by KDM2B. Therefore, in the context of the work presented in this report, KDM2B functions as a transcriptional activator, and not as a transcriptional repressor, as it has been observed in other contexts [4, 8, 15]. Consistent with earlier observations showing that transcriptional repression by KDM2B selectively depends on the functional cooperation between KDM2B and EZH2 [4, 8], the knockdown of EZH2 had no effect on the transcription activator function of KDM2B. However, disruption of the variant non-canonical polycomb complex ncPRC1.1, which is targeted to DNA by KDM2B, one of its components [7, 45], abolished the KDM2B transcription activator function, suggesting that KDM2B activates transcription in the context of ncPRC1.1. ATAC-Seq and ChIP-Seq experiments in control and KDM2B knockdown cells, along with bioinformatics analyses of a large dataset of 465 TNBC patients (360 with RNA-Seq data) (FUSCC cohort) [46, 47] provided evidence suggesting that KDM2B regulates not only the expression, but also the transcriptional activity of MYC. Overall, the data in this report show that KDM2B plays an important role in the global regulation of cellular metabolism. In the SGOC pathway, which is the focus of this report, KDM2B regulates the SGOC enzyme-encoding gene activators, ATF4 and MYC, and functions as a master regulator of the pathway.

## Results

### The knockdown of KDM2B promotes oxidative stress and renders cancer cells sensitive to APR-246 and deubiquitinase inhibitors

KDM2B is overexpressed and promotes tumorigenesis of several types of tumors. In human breast cancer, it is overexpressed selectively in basal-like TNBC (**Fig. 1A**), which is the focus of this report. Our earlier studies had shown that overexpression of KDM2B in MEFs renders them resistant to oxidative stress. The same studies showed that this phenotype was due to the KDM2B-dependent regulation of antioxidant mechanisms [15]. To determine whether KDM2B is also required for the activity of antioxidant mechanisms in basal-like breast cancer, we knocked down KDM2B in MDA-MB-231 and MDA-MB-468 cells [48] and we examined the levels of ROS in empty vector (EV)-control and shKDM2B-transduced cells (**Fig. 1B**), before and after treatment with TBHP. This showed that the knockdown (KD) of KDM2B indeed results in an increase of the intracellular levels of ROS, both before and after treatment with TBHP (**Fig. 1C**). Consistent with this observation, the KD of KDM2B induces apoptosis as evidenced by increased Annexin-V staining **(Fig. S2A,B).** Additionally, shKDM2B-transduced cells were sensitized to the ROS inducer Piperlongumine (PL), as evidenced by the fact that they exhibited reduced viability when treated with the drug for 48 hours (**Fig. 1D, S1).**

**Figure 1.**
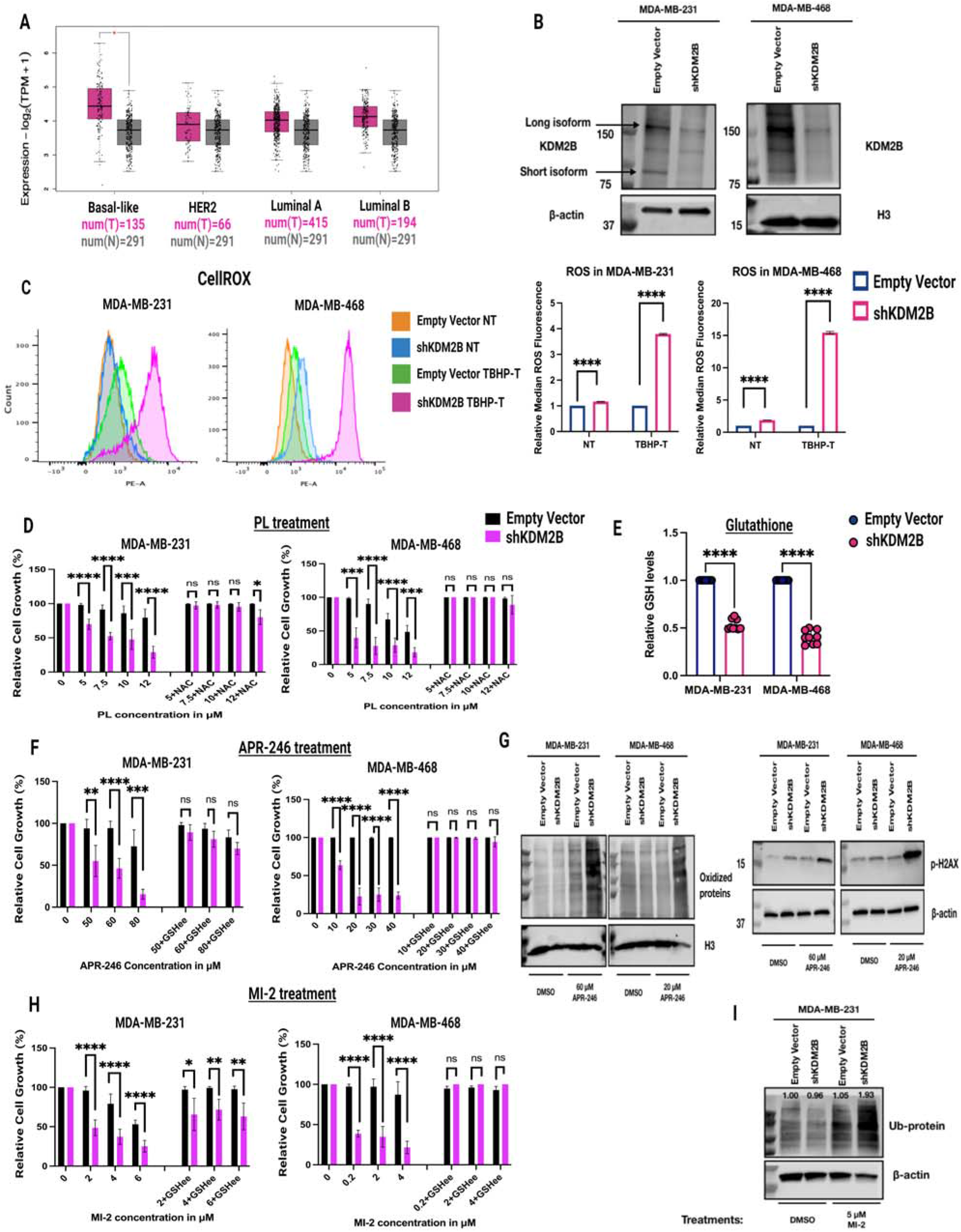
The knockdown of KDM2B promotes oxidative stress and renders cancer cells sensitive to APR-246 and deubiquitinase inhibitors. **A)** Boxplots showing the expression of KDM2B in basal-like, Her2-positive, Luminal A, and Luminal B breast cancer and comparing the expression in tumor and normal samples. TCGA data, downloaded from Gepia2 (Log2FC=0.6, p-value <0.05). Statistical significance was determined by one-way ANOVA. **B)** Immunoblots of Empty Vector and shKDM2B-transduced MDA-MB-231 (left panel) and MDA-MB-468 (right panel) cells, were probed with the anti-KDM2B antibody. Arrows indicate the long and short isoform of KDM2B. β-actin and Histone H3 were used as loading controls. **C)** ROS levels in Empty vector and shKDM2B-transduced MDA-MB-231 and MDA-MB-468 cells. <u>Left panel:</u> ROS levels, were measured by flow cytometry in cells pretreated with the oxidative stress indicator CellROX (orange fluorescence). NT= non-treated, TBHP T= treated with 100 nM tert-Butyl hydroperoxide (TBHP) for 3h. <u>Right panel</u>: Quantification of the flowcytometry histograms in the upper panel, presented as Median Fluorescence Intensity (MFI). Data are from three biological replicates, and they are presented as mean ± SD. Statistical significance was determined with the unpaired two-tailed t-test. **The knockdown of KDM2B sensitizes the cells to the ROS inducer Piperlogumine (PL), the pro-oxidant APR-246, and the Deubiquitinase (DUB) inhibitor MI-2.** **D)** The KD of KDM2B sensitizes cells to PL. Similar numbers of Empty vector and shKDM2B-transduced MDA-MB-231 and MDA-MB-468 cells were treated with PL or DMSO, starting at 24 hours after seeding. Parallel cultures were also treated with 5mM of the ROS scavenger, NAC. Cell proliferation was monitored in the Incucyte live-cell analysis system. Figure shows the cell confluency readout of the Incucyte (a measure of relative cell number) at 48-hours from the start of the treatment. Relative cell numbers are presented as a percentage of the cell growth in the treated conditions compared to the cell growth in the non-treated (DMSO) conditions (100% cell growth). Figure shows the mean value ± SD of the combined results of two independent experiments, each of which included three biological replicates (n=6). Statistical significance was determined with the unpaired two-tailed t-test. **E)** The KD of KDM2B lowers the levels of reduced Glutathione (GSH) in MDA-MB-231 and MDA-MB-468 cells. Figure shows the mean value ± SD of the combined results of three independent experiments, each of which included three biological replicates (n=9). Statistical significance was determined with the unpaired two-tailed t-test. **F)** The KD of KDM2B sensitizes cells to APR-246. Sensitivity of Empty Vector and shKDM2B cells to APR-246 was done, as described for PL sensitivity in the upper panel. The phenotype of APR-246 treated cells was rescued with 5 mM of GSHee (Glutathione ethyl-ester). Figure shows the mean value ± SD of the combined results of two independent experiments, each of which included three biological replicates (n=6). Statistical significance was determined with the unpaired two-tailed t-test. **G)** <u>Left panel</u>. Oxyblot of Empty Vector and shKDM2B-transduced MDA-MB-231 and MDA-MB-468 cells, treated with DMSO, or APR-246 (60 μM for MDA-MB-231 and 20 μM for MDA-MB-468) for 24 hours. The loading control was histone H3. <u>Right panel</u>. Immunoblots of Empty Vector and shKDM2B-transduced MDA-MB-231 and MDA-MB-468 cells, treated with DMSO, or APR-246 (60 μM for MDA-MB-231 and 20 μM for MDA-MB-468) for 24 hours, were probed with anti-phospho-H2AX (gamma H2AX). Loading control was β-actin. **H)** The KD of KDM2B sensitizes cells to MI-2. Sensitivity of Empty Vector and shKDM2B cells to MI-2 was done, as described in the upper and middle panels. The phenotype MI-2-treated cells was again rescued with 5 mM of GSHee (Glutathione ethyl-ester). Figure shows the mean value ± SD of the combined results of two independent experiments, each of which included three biological replicates (n=6). Statistical significance was determined with the unpaired two-tailed t-test. **I)** Immunoblot of Empty Vector and shKDM2B-transduced MDA-MB-231 cells, treated with 5 μM MI-2 or DMSO, was probed with the anti-Ub antibody. β-actin was used as loading control.

The significantly higher levels of ROS in shKDM2B cells treated with TBHP also suggested that in agreement with our earlier data, the shKDM2B-associated accumulation of ROS was due to a defect in ROS scavenging, and this was further supported by the observation that NAC [49], rescued the shKDM2B-induced oxidative stress phenotype (**Fig. 1D**). Given that the most abundant antioxidant is GSH, we measured the cellular levels of reduced GSH [50] and we found that its levels were significantly decreased in the KDM2B KD cells (**Fig. 1E**).

Tumor cells with low levels of GSH, due to the low abundance of the glutamate-cystine antiporter xCT/SLC7A11, tend to be sensitive to Eprenetapopt (APR-246) [51, 52], a small molecule, which in the cellular environment is converted to methylene quinuclidinone (MQ). The latter binds thiol groups, including GSH, and perturbs the antioxidant balance of the cells. The pro-oxidant and cytotoxic activities of APR-246 have been attributed to the inhibition and/or depletion of GSH and to the inhibition of thioredoxin [51]. APR246 is also known to selectively target tumor cells with mutations in p53, and this may also be due at least in part to the low abundance of GSH in such tumors [52]. Based on these considerations we treated control and shKDM2B-transduced MDA-MB-231 and MDA-MB-468 cells with APR-246 and we observed that the KD of KDM2B renders them sensitive to the drug (**Fig. 1F, S1,S2C)**. Moreover, APR-246 treatment led to increased levels of oxidized proteins and γ-H2AX in shKDM2B cells (**Fig. 1G**). In addition, the enhanced toxicity of APR-246 in the shKDM2B cells was due to the low abundance of GSH, as it was reversed by GSH supplementation (**Fig. 1F**).

Earlier studies had shown that TNBC cells tend to be resistant to GSH depletion by activating enzyme systems that deubiquitinate proteins that cause ER and proteotoxic stress. As a result, DUB inhibition, in combination with GSH depletion promotes proteotoxic stress and cell death [53]. Based on this information we examined the sensitivity of control and shKDM2B-transduced cells to the DUB inhibitor MI-2 (MALT-1) [53] and we found that the KD cells are indeed selectively sensitive to this treatment as evidenced by the monitoring of cell growth (**Fig. 1H, S1)** and PARP cleavage **(Fig. S2D).** The activity of MI-2 in this experiment was confirmed by the increased protein ubiquitination in the treated cells (**Fig. 1I**). More importantly, their enhanced sensitivity was again dependent on the low levels of GSH, as it was reversed by GSH supplementation (**Fig. 1H**).

Potential mechanisms for the regulation of GSH levels by KDM2B include the expression and activity of the antiporter xCT/SLC7A11 [52], the expression of GLS, the expression and activity of the rate limiting GCLC and GCLM, and the expression and activity of GSS [50]. The relative ratio of reduced and oxidized Glutathione (GSH/GS-SG) also depends on GPX4 and the activity of GSR [50]. Western blotting addressing the expression of all these molecules however, showed no differences between the control and shKDM2B cells **(Fig. S3A-C).**

### Transcriptomic, proteomic and metabolomic profiling reveals major shifts in metabolism in shKDM2B-transduced MDA-MB-231 cells

Given that the expression of the xCT/SLC7A11 antiporter and the enzymes regulating the synthesis of GSH was not affected by the knockdown of KDM2B, we hypothesized that KDM2B may regulate the biosynthesis and degradation of the GSH building blocks, glutamate, cysteine and glycine. To address this question in an unbiased way, we employed RNA-Seq, TMT quantitative proteomics, and LC-MS untargeted metabolomics, to compare the transcriptomes, proteomes, and metabolomes of control and shKDM2B-transduced MDA-MB-231 cells.

Analysis of the transcriptomic and proteomic data revealed major shifts in intermediary metabolism. Ridge plot of the differentially expressed genes focusing on the GO domain “Biological Process” (BP) revealed that genes downregulated in the KDM2B knockdown cells are enriched for the GO terms “alpha-amino acid biosynthetic process” and “cellular amino acid biosynthetic process” or the related terms “cellular amino acid metabolic process” and “alpha amino acid metabolic processs” (**Fig. 2A, S4A).** Analysis of the same data with MITHril, a neural network-based topological pathway tool, was consistent with the Ridge plot analysis by showing that the top enriched term was “Biosynthesis of amino acids”. Importantly many additional terms associated with amino acid metabolism were also enriched (**Fig. 2B**). GSEA confirmed the significant enrichment of “cellular amino acid metabolic processes” **(Fig. S4B).** The differential expression of genes involved in these pathways is presented in the heatmaps in figure 2C (**Fig. 2C, S4C,E).** Analysis of the TMT-proteomics data showed a significant overlap with the transcriptomic data (**Fig. 2D, S4D,F).**

**Figure 2.**
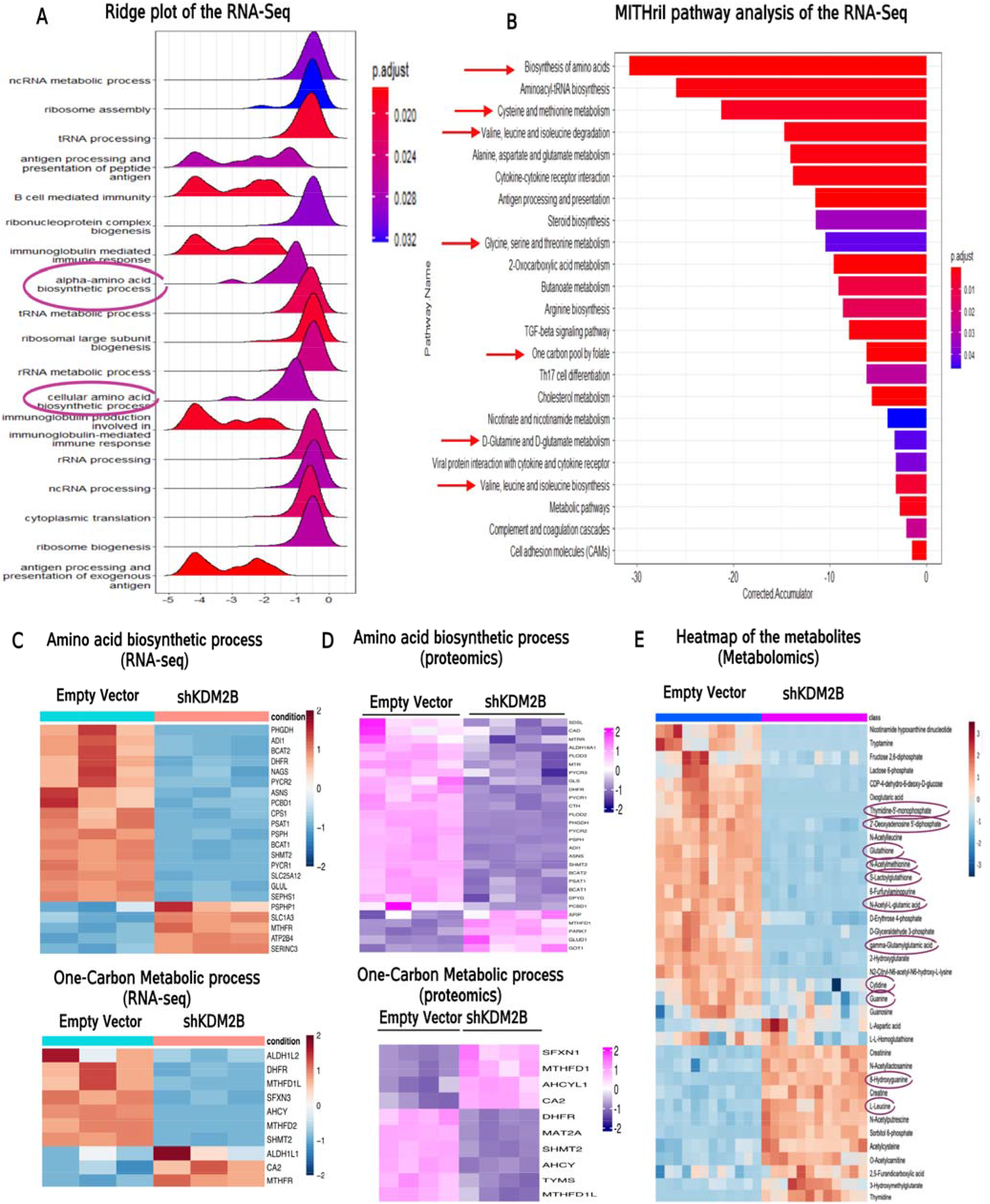
Transcriptomic, proteomic and metabolomic profiling reveals major shifts in metabolism. **A)** Comparison of the transcriptomes of Empty Vector and shKDM2B-transduced MDA-MB-231 cells. RNA-seq was performed on three biological replicates. Ridge plot analysis of the RNA-seq data, based on genes with statistically significant log2FC and the GO Domain “Biological Process”. Alpha-amino acid and Cellular amino acid biosynthetic processes were among the statistically significant downregulated processes in shKDM2B cells. **B)** Analysis of the RNA-seq data using MITHril, a neural network-based topological pathway tool utilizing the information in the KEGG metabolic pathways. This also shows downregulation of “biosynthesis of amino acids” in shKDM2B-transduced cells. The related Cysteine and methionine metabolism, Valine leucine and isoleucine biosynthesis and degradation, Glycine, serine, and threonine metabolism, one carbon metabolism and D-glutamine and D-Glutamate metabolism pathways, were also among the pathways, exhibiting statistically significant downregulation in the shKDM2B cells. **C)** Heatmaps of the genes in GO “amino acid biosynthetic process” and “one-carbon metabolic process” that are differentially expressed in Empty vector and shKDM2B cells. **D)** Heatmaps of the proteins encoded by genes in GO “amino acid biosynthetic process” and “one-carbon metabolic process” that are differentially expressed in Empty vector and shKDM2B cells. TMT Proteomics data derived from the analysis of four biological replicates. **E)** Heatmap of differentially abundant metabolites, detected by untargeted metabolomics in Empty vector and shKDM2B-transduced MDA-MB-231 cells (four biological replicates per cell type). Significant differences in the abundance of metabolites regulated by enzymes in GO “amino acid biosynthetic process” were observed between control and KDM2B knockdown cells. GSH was also reduced and 2-hydroxyguanine, a marker of oxidative DNA damage, was elevated.

The abundance of the GSH building blocks, cysteine, glutamate, and glycine, depends on the activity of the SGOC pathway, the transsulfuration pathway [23, 25] and the availability of glutamate [50]. The latter is a non-essential amino acid which may be synthesized from glutamine via deamination reactions catalyzed by GLS or ASNS, or from BCAAs via pathways regulated by the BCAT1/2 [54, 55]. Our analyses of the transcriptomic and proteomic data described above, provided evidence that the SGOC, transsulfuration and BCAA-dependent glutamate synthesis pathways are among the most downregulated metabolic pathways in shKDM2B-transduced MDA-MB-231 cells. Additionally, whereas the levels of GLS were not affected by the knockdown of KDM2B **(Fig. S3C),** the expression of ASNS was significantly lower in shKDM2B-transduced cells (**Fig. 2C, D).**

Importantly, the expression of a subset of the KDM2B-regulated SGOC enzymes is higher in the basal-like TNBC as opposed to the other breast cancer subtypes, in agreement with earlier observations [31, 32, 34, 36] **(Fig. S5A).** Additionally, analysis of the transcriptomic data in two cohorts of TNBC patients [46, 47], showed that the expression of these enzymes correlates significantly with the expression of KDM2B **(Fig. S5B).**

LC-MS metabolomics detected 36 unambiguously-identified metabolites. Of those, 21 were downregulated and 13 were upregulated in shKDM2B cells (**Fig. 2E**). The metabolomics data confirmed that GSH, and the related metabolites S-lactoglutathione, N-acetyl-glutamic acid, and gamma-glutamyl-glutamic acid were downregulated in the shKDM2B cells. Additionally, the levels of the glutamate precursor L-leucine were increased as expected from the downregulation of BCAT1/2. Consistent with the downregulation of GSH, the abundance of 8-OH-Gua, a marker of oxidative DNA damage, was increased. The increase in the abundance of the antioxidant acetyl-cysteine, which was also observed, was probably a compensatory survival mechanism [56]. The combined transcriptomic, proteomic and metabolomic data were consistent with our hypothesis that the effect of KDM2B on the abundance of GSH is due to the KDM2B-dependent regulation of the availability of the GSH building blocks.

The observed decrease in the abundance of both purines (Guanine, and 2-deoxyadenosine-5-diphosphate), pyrimidines (Thymidine-5’-monophosphate, and Cytidine) and N-acetyl-methionine was consistent with the downregulation of the 1C metabolism pathway. In addition, it was consistent with the apparent downregulation of glycolysis and the non-oxidative branch of PPP (decrease in the abundance of Fructose-2,6-Biphosphate, Glyceraldehyde-3-phosphate and D-Erythrose-4-phosphate). The discordant and unexpected increase in the abundance of thymidine in the shKDM2B cells suggests that although the biosynthesis of thymidine may be intact, its utilization may be defective. Consistent with this interpretation is the observed significant downregulation of thymidine-5-monophosphate (**Fig. 2E**).

### KDM2B regulates SGOC and glutamate metabolism in basal-like breast cancer cells and its knockdown impairs cell proliferation in culture and tumor growth in mice

The preceding data provided evidence that the knockdown of KDM2B in MDA-MB-231 cells reduces the activity of SGOC and glutamate biosynthesis pathways which control the biosynthesis of glutathione. The role of these enzymes in the regulation of these pathways is illustrated in the pathway diagram in Fig. 3A (**Fig. 3A**). To validate the transcriptomic and proteomic data supporting the role of KDM2B in SGOC and glutamate metabolism, we selected a set of enzymes, some of which play a rate limiting role in these pathways, and we examined their expression by qRT-PCR and western blotting in control and shKDM2B cells (**Fig. 3B, C)**.

**Figure 3.**
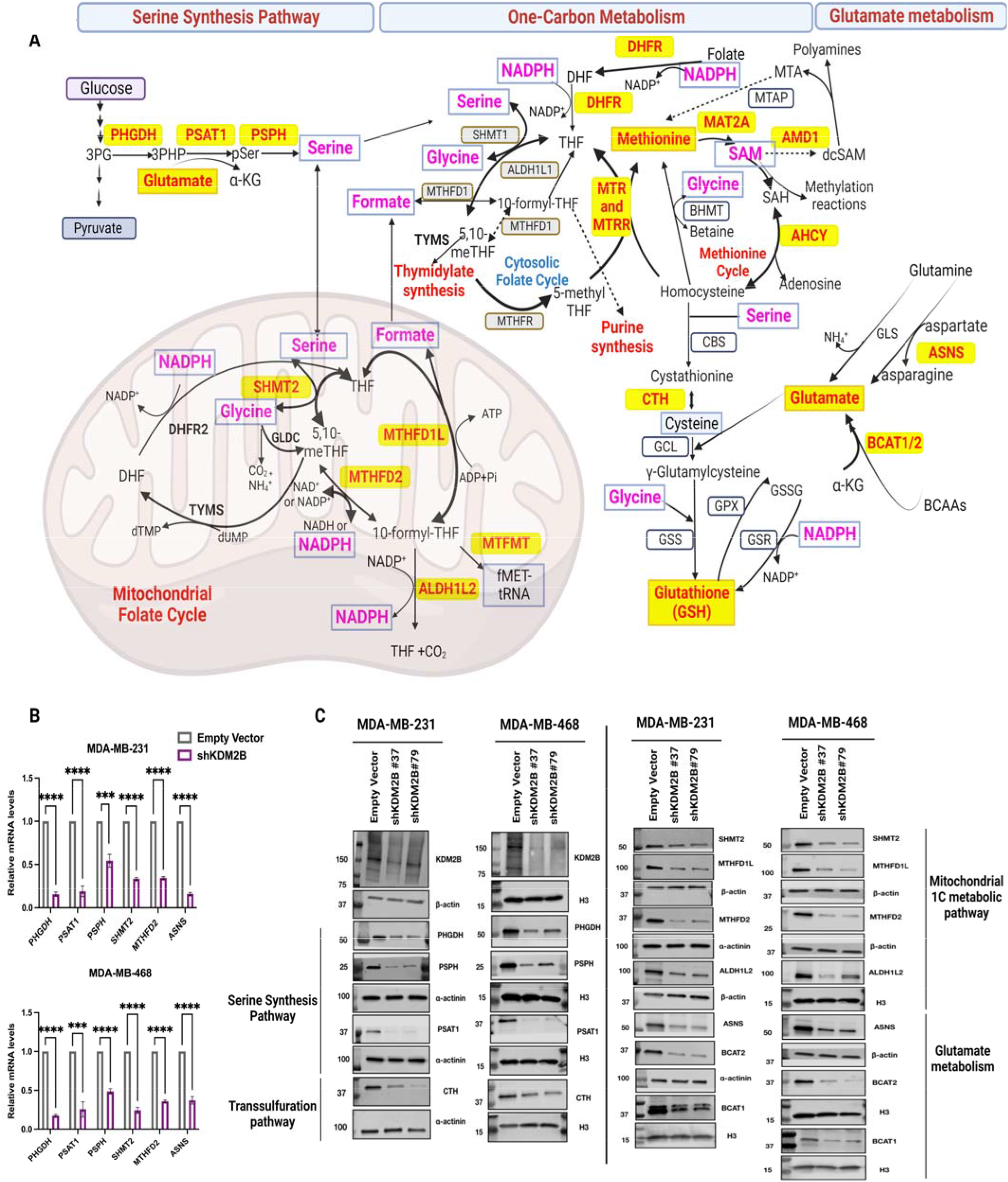
KDM2B regulates the SGOC, transsulfuration and glutamate metabolism pathways in basal-like breast cancer cells. **A)** Schematic diagram of the SSP, 1C metabolism, glutamate metabolism and GSH synthesis pathways. Serine can be taken up from the environment or synthesized from the glycolytic intermediate 3-phosphoglycerate (3-PG). The first, rate limiting step in <u>serine biosynthesis</u> is under the control of phosphoglycerate dehydrogenase (PHGDH), which catalyzes the oxidation of 3-PG to 3-phosphohydroxypyruvate (3-PHP). 3-PHP is then converted into 3-phosphoserine (3-PS) via a glutamate-dependent transamination reaction, which is catalyzed by phosphoserine aminotransferase 1 (PSAT1) and produces a-ketoglutarate (a-KG), in addition to 3-PS. In the next and final step phosphoserine phosphatase (PSPH) catalyzes the hydrolysis of 3-PS to produce serine. <u>1C metabolism</u> is organized around two interconnected methylation cycles, which operate in the mitochondria and the cytoplasm. The major donor of 1C units is serine, which donates a methyl group to tetrahydrofolate (THF), producing 5,10-methylene-THF (5,10 me-THF) and glycine, in a reaction catalyzed by serine hydroxymethyltransferase (SHMT1 in the cytosol, and SHMT2 in the mitochondria). Oxidation of 5,10-me-THF by methylenetetrahydrofolate dehydrogenase (NADP+-dependent MTHFD1 in the cytosol and NAD, or NADP+-dependent MTHFD2 in the mitochondria), produces 10-formyl-THF and generates NADH, or the reducing agent, NADPH. 10-formyl-THF releases formate via a reaction catalyzed by the NADP+ dependent enzymes methylenetetrahydrofolate dehydrogenase 1 like (MTHFD1L) in the mitochondria and MTHFD1 in the cytosol, and the released formate diffuses between subcellular compartments and is utilized for the biosynthesis of purine bases. Moreover, 10-formyl-THF can be used to generate 5,10-me-THF in both the mitochondria and the cytosol, and 5,10-me-THF can function as a methyl donor to produce dTMP from dUMP, in a reaction catalyzed by thymidylate synthase (TYMS). In the mitochondria, 10-formyl-THF is also used as a formyl donor to generate formylmethionyl-tRNA (f-Met tRNA), via a reaction catalyzed by methionyl-tRNA formyltransferase (MTFMT). Additionally, THF can be recovered from 10-formyl-THF, via a reaction which releases CO2 and is catalyzed by Aldehyde Dehydrogenase 1 Family Member L2 (ALDH1L2). <u>Methionine cycle</u>. 5,10-meyhyl-THF reductase (MTHFR) catalyzes the conversion of 5,10-meTHF to 5-methyl-THF in the cytosol. The latter functions as a methyl donor for the methylation of homocysteine to methionine, in a reaction catalyzed by Methionine Synthase (MTR), and its Vitamin B12-dependent regulator Methionine Synthase Reductase (MTRR). An alternate pathway for the remethylation of homocysteine, utilizes betaine as the methyl-donor and is catalyzed by Betaine-Homocysteine-Methyltransferase, (BHMT) and its co-factor Vitamin B12. Adenylation of methionine by methionineadenosyltransferase 2A (MAT2A), produces S-adenosylmethionine (SAM), which serves as the methyl donor in a large number of methylation reactions, converting SAM to S-adenosylhomocysteine (SAH). The latter is hydrolyzed by Adenosylhomocysteinase (AHCY), closing the methylation cycle by producing adenosine and homocysteine. SAM, may also be decarboxylated by Adenosylmethionine decarboxylase 1 (AMD1), and decarboxylated SAM (dcSAM) is used for polyamine synthesis. Methyl-thio-adenosine (MTA) produced in the course of polyamine synthesis, enters a salvage pathway which depends on Methyl-Thio-Adenosine-Phosphorylase (MTAP) to regenerate methionine. <u>Transsulfuration pathway</u>. Coupled to the methionine cycle is transsulfuration, a pathway defined by the transfer of sulfur from methionine to serine via homocysteine, which is the immediate sulfur donor in the pathway. The first step is the condensation of homocysteine and serine to form cystathionine, in a reaction catalyzed by cystathionine β-synthase (CBS). In the last step of the transsulfuration pathway, cystathionine γ-lyase (cystathionase, CTH) catalyzes the cleavage of Cystathionine to generate cysteine and α-keto-butyrate. <u>Glutamate metabolism and GSH biosynthesis.</u> The first rate-limiting step in GSH biosynthesis is the ATP-dependent ligation of Cysteine to Glutamate, which produces γ-glutamylcysteine. This reaction is catalyzed by Glutamate-Cysteine Ligase (GCL), an enzyme consisting of two subunits, a heavy catalytic subunit (GCLC) and a light regulatory subunit (GCLM). Glutamate, which feeds into this reaction is produced by Glutamine, or Branched Chain Amino Acids (BCAAs, Leucine, Isoleucine and Valine), via reactions that are catalyzed by Glutaminase (GLS), Asparagine Synthetase (ASNS), or Branched Chain Amino Acid Transaminase (BCAT), as indicated. Glutathione Synthetase (GSS) catalyzes the condensation of γ-glutamylcysteine and glycine, which is the last step in GSH biosynthesis. The antioxidant function of GSH is catalyzed by Glutathione peroxidases (GPX) and the oxidized Glutathione (GSSG) generated by GPX-dependent reactions, is reduced back to GSH by Glutathione Reductase (GSR) using NADPH as a cofactor (Yang and Vousden 2016; Dekhne et al., 2020; Geeraerts et al., 2021; Lieu et al., 2020). Metabolites and enzymes whose abundance/expression is downregulated by the knockdown of KDM2B (based on our multi-omic analysis) are shown in red and they are framed in yellow boxes. Metabolites whose abundance is lower in shKDM2B cells (based on our in *vitro* biochemical assays) are shown in magenta and they are framed in blue boxes. **B)** Relative m-RNA levels of enzymes in the SGOC and glutamate metabolism pathways in Empty Vector and shKDM2B-transduced MDA-MB-231 and MDA-MB-468 cells, were measured by qRT-PCR. The control was β-actin. Data from three biological replicates, presented as mean ± SD, (n=3). Statistical significance was determined with the unpaired two-tailed t-test. **C)** Representative immunoblots of KDM2B and multiple metabolic enzymes involved in SGOCP (Serine Glycine and Mitochondrial 1C metabolism pathway), and the transsulfuration and glutamate metabolism pathways in Empty Vector and shKDM2B MDA-MB-231 and MDA-MB-468 cells. The loading controls were histone H3, β-actin and α-actinin, as indicated. KDM2B was knocked down with two different shRNAs (#37, #79).

As mentioned in the preceding section, our metabolomics data were consistent with the transcriptome and proteome-based evidence indicating that the SGOC and glutamate biosynthesis pathways are regulated by KDM2B. However, due to the low depth of the metabolomics data, only few critical metabolites had been detected. To address this problem, we examined the abundance of serine, glycine, formate, NADPH, glutamate and SAM in lysates of control and shKDM2B-transduced MDA-MB-231 and MDA-MB-468 cells, using in vitro biochemical assays. The results confirmed that the knockdown of KDM2B led to a significant downregulation of the abundance of all six metabolites and provided additional support for the positive regulation of these pathways by KDM2B (**Fig. 4A**).

**Figure 4.**
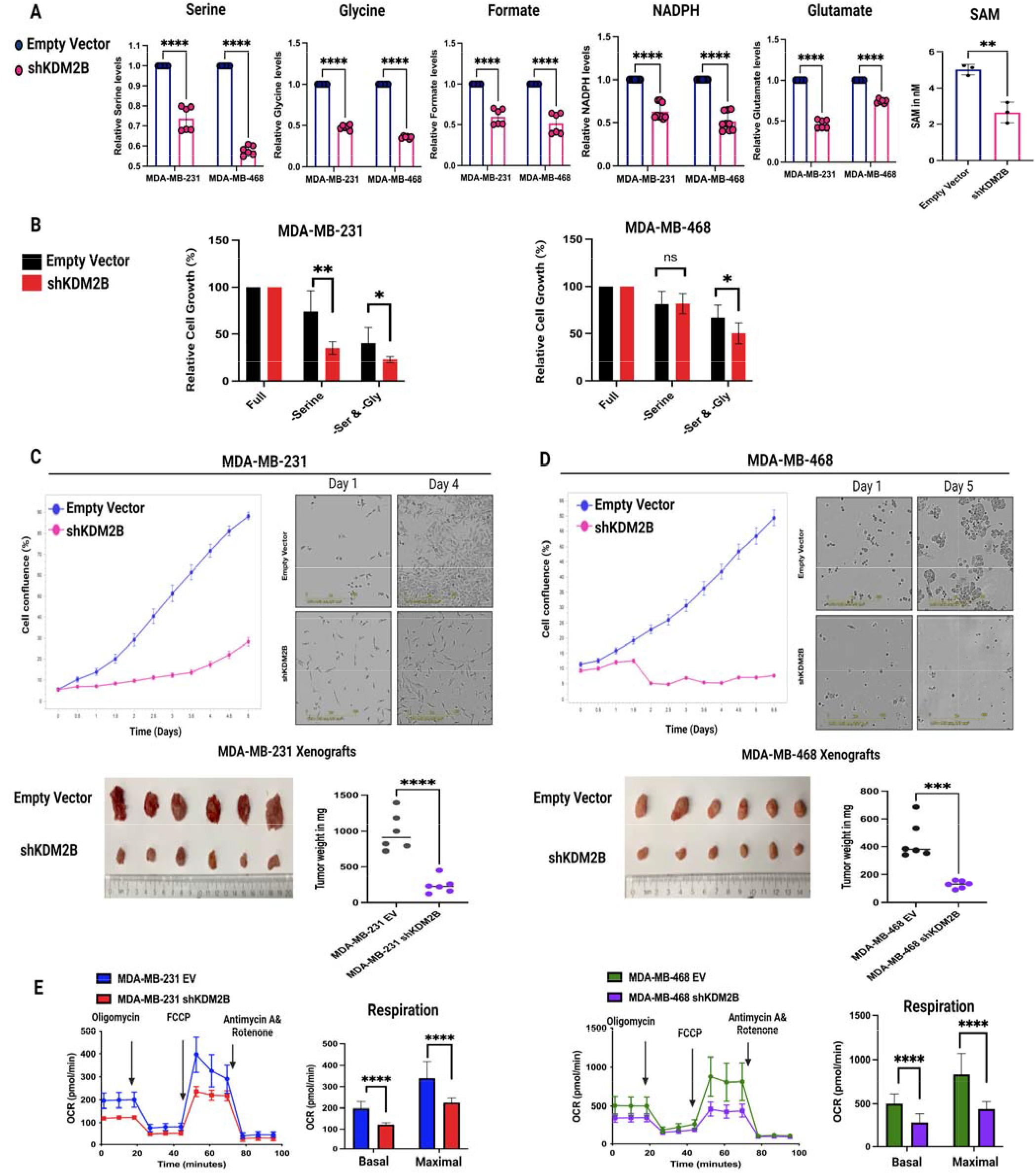
KDM2B regulates the abundance of SGOCP metabolites and the rate of mitochondrial respiration, and its knockdown impairs cell proliferation in culture and tumor growth in mice. **A)** The knockdown of KDM2B in MDA-MB-231 and MDA-MB-468 cells lowers the abundance of serine, glycine (inputs of the 1C-metabolism), formate, NADPH, SAM (outputs of the 1C-metabolism) and glutamate. Data from two independent experiments with three biological replicates each (n=6) are presented as the mean ± SD. SAM levels were measured only in MDA-MB-231 cells using three replicates, n=3. Statistical significance was determined with the unpaired two-tailed t-test. **B)** Relative cell numbers of Empty Vector and shKDM2B-transduced MDA-MB-231 (left panel) and MDA-MB-468 cells (right panel), grown in complete (Full), serine deficient (-Serine) or serine and glycine deficient (-Ser&Gly) media. Relative cell numbers were based on Incucyte images, which were captured on day 5 from the day the cells were seeded. Relative cell numbers are presented as a percentage of the cell growth in the deficient media compared to the cell growth in full media (100% cell growth). Figure shows the mean value ± SD of the combined results of two independent experiments, each of which included three biological replicates (n=6). Statistical significance was determined with the unpaired two-tailed t-test. **C-D**). Proliferation of Empty Vector and shKDM2B-transduced MDA-MB-231 and MDA-MB-468 cells in culture and growth of tumor xenografts initiated by subcutaneous injection of the same cells in the flanks of immunocompromised mice. <u>Upper panels</u>: Cell proliferation curves in the left upper panels of C and D describe the rates of increasing cell confluence of cultured cells, which was monitored with the Incucyte live-cell analysis system. Data were from triplicate cultures (n=3). Representative images of the same cultures at day 1 and day 4 (MDA-MB-231 cells) or day 5 (MDA-MB-468 cells) are shown in the right upper panels of C and D. <u>Lower</u> <u>panels</u>: Size and weight of both the MDA-MB-231 and MDA-MB-468 tumor xenografts were reduced by the knockdown of KDM2B. The weight of the xenografts is presented as the mean ± SD of six xenografts (n=6) from each cell line. Statistical significance was determined with the unpaired two-tailed t-test. **E)** Oxygen consumption rate (OCR) of Empty Vector and shKDM2B-transduced MDA-MB-231 and MDA-MB-468 cells was measured with the Seahorse XF Cell Mito Stress Test kit. Complex V inhibitor oligomycin, the uncoupler carbonyl cyanide 4-(trifluoromethoxy)phenylhydrazone (FCCP), and the complex I and III inhibitors Antimycin A and Rotenone were added sequentially at the indicated time points. OCR data after normalization based on the cell number measured with the incucyte live-cell analysis system. Data are presented as the mean ± SD of six biological replicates (n=6). Statistical significance was determined with the unpaired two-tailed t-test.

Earlier studies had shown that inhibition of PHGDH, and MTHFD2, significantly impairs cell proliferation [21, 31, 36, 57]. Experiments in MDA-MB-231 and MDA-MB-468 cells presented this report, confirmed these observations. In addition, they showed that the combination of PHGDH, and MTHFD2 inhibition with the KDM2B knockdown, which downregulates these enzymes, almost completely blocks cell proliferation **(Fig. S6A,B)**. The importance of serine and glycine in this phenotype was confirmed by monitoring the proliferation of control and shKDM2B-transduced MDA-MB-231 and MDA-MB-468 cells, following serine, or serine-glycine deprivation (**Fig. 4B**). MDA-MB-468 cells carry an amplified *PHGDH* locus [31] and they are less sensitive to serine and serine-glycine deprivation, as well as to the combination of amino acid deprivation and KDM2B knockdown, as expected (**Fig. 4B**).

Earlier studies [7, 9, 10, 13, 14], and data presented in this report (**Fig. 4C,D**) confirmed that shKDM2B inhibits the proliferation of cancer cells, both in culture and in mouse xenograft models. The experiment in figures 4C and 4D also showed that MDA-MB-468 cells, which carry the amplified *PHGDH* locus, and are therefore addicted to the SGOC pathway, are significantly more sensitive to the knockdown of KDM2B. These data combined, strongly suggest that SGOC inhibition contributes to the inhibition of cell proliferation by shKDM2B. This conclusion is supported by the results of the mitochondrial respiration experiment in figure 4E, which shows that the effect of the KDM2B knockdown on OCR is more robust in the SGOC-addicted MDA-MB-468 cells (**Fig. 4E**). Mitochondrial respiration provides energy for cell proliferation and depends on serine metabolism [58, 59], which is regulated by KDM2B. Therefore, its inhibition by shKDM2B, which was more pronounced in the SGOC addicted MDA-MB-468 cells, strongly supports the conclusion that SGOC inhibition is a critical contributor to the reduction of cell proliferation by shKDM2B.

### KDM2B regulates a network of epigenetic and transcription factors, which control the expression of enzymes in the SGOC, glutamate and GSH metabolism pathways

The main transcription factors regulating the expression of the enzymes that control the metabolic pathways in this report are ATF4 and MYC [22, 24, 41, 42, 43, 44, 60, 61] whose expression, based on our transcriptomic data, is regulated positively by KDM2B. Although multiple studies to-date have shown that ATF4 is regulated at the level of translation [60, 62, 63], the downregulation of its mRNA in shKDM2B-transduced cells suggests that KDM2B may regulate it, similarly to MYC, at the level of transcription (**Fig. 5A,B**). We should add that the transcriptional regulation of MYC by KDM2B was also supported by exon-intron split analysis (EISA) of our RNA-Seq data, which showed that the number of intronic reads of MYC and some of the SGOC target genes was significantly reduced in the shKDM2B cells (**Fig. 5C**).

**Figure 5.**
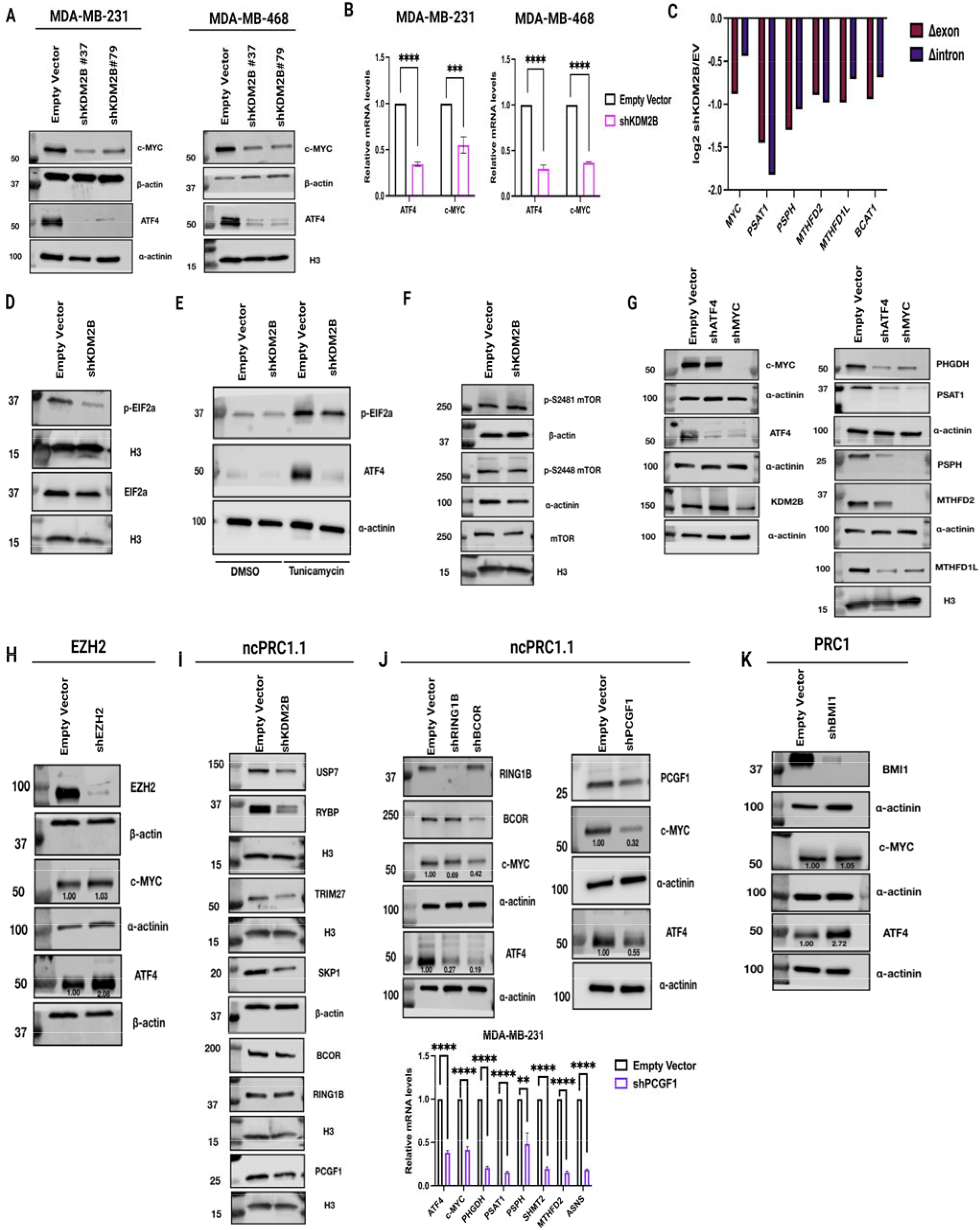
KDM2B regulates a network of transcription and epigenetic factors, which control the expression of enzymes in the SGOC and glutamate metabolism pathways. **A)** Immunoblots of Empty Vector and shKDM2B-transduced MDA-MB-231 and MDA-MB-468 cells, probed with anti-MYC or anti-ATF4 antibodies. KDM2B was knocked down with two different shRNAs (#37, #79). Histone H3, β-actin and α-actinin were used as loading controls, as indicated. **B)** Relative m-RNA levels of MYC and ATF4 in Empty Vector and shKDM2B-transduced MDA-MB-231 and MDA-MB-468 cells were measured by qRT-PCR. KDM2B was knocked down with shRNA #79. The control was β-actin. Mean ± SD of three independent biological replicates (n=3). Statistical significance was determined with the unpaired two-tailed t-test. **C)** Exon-Intron Split Analysis (EISA) shows the log2 ratio of the number of exonic (Δexon) and intronic (Δintron) reads of MYC and the genes encoding several KDM2B-regulated metabolic enzymes, in shKDM2B/Empty Vector MDA-MB-231 cells. **D)** Representative immunoblot of Empty Vector and shKDM2B-transduced MDA-MB-231 cells, probed with the anti-p-EIF2a and EIF2α antibodies. KDM2B was knocked down with shRNA #79. The loading control was Histone H3. **E)** Representative immunoblots of Empty Vector and shKDM2B-transduced MDA-MB-231 cells, treated with the ER stress inducer tunicamycin (1μg/μl) or DMSO for 16 hours, and probed with the anti-p-EIF2a or ATF4 antibodies. KDM2B was knocked down with shRNA #79. The loading control was α-Actinin. **F)** Representative immunoblots of Empty Vector and shKDM2B-transduced MDA-MB-231 cells, were probed with the anti-p-S2448, p-S2481 and total mTOR antibodies. KDM2B was knocked down with shRNA #79. The loading controls were β-actin, α-actinin, or Histone H3, as indicated. **G)** Representative immunoblots of Empty Vector, shATF4, or shMYC-transduced MDA-MB-231 cells, were probed with anti-MYC, ant-ATF4, and anti-KDM2B antibodies, or with antibodies to several metabolic enzymes encoded by KDM2B-regulated genes (PHGDH, PSAT1, PSPH, MTHFD2 and MTHFD1L). The loading controls were α-Actinin and histone H3, as indicated. **H)** Representative immunoblots of Empty Vector and shEZH2-transduced MDA-MB-231 cells, probed with anti-EZH2 anti-MYC, and anti-ATF4 antibodies. The loading controls were β-actin and α-actinin. **I)** Representative immunoblots of Empty Vector and shKDM2B-transduced MDA-MB-231 cells, probed with antibodies to USP7, RYBP, TRIM27, SKP1, BCOR, RING1B and PCGF1. KDM2B was knocked down with shRNA #79. The loading controls were histone H3 and β-actin. **J)** <u>Upper panels:</u> Representative immunoblots of Empty Vector and shRING1B, shBCOR and shPCGF1-transduced MDA-MB-231 cells, were probed with antibodies to RING1B, BCOR, PCGF1, c-MYC, or ATF4, as indicated. The loading control was α-actinin. <u>Lower panel:</u> Relative m-RNA levels of ATF4, c-MYC, PHGDH, PSAT1, PSPH, SHMT2, MTHFD2 and ASNS in Empty Vector and shPCGF1-transduced MDA-MB-231 cells, measured by qRT-PCR. The loading control was β-actin. Mean ± SD of three replicates (n=3). Statistical significance was determined with the unpaired two-tailed t-test. **K)** Representative immunoblots of Empty Vector and shBMI1-transduced MDA-MB-231 cells, probed with antibodies to BMI1, c-MYC and ATF4, as indicated. The loading control was α-actinin.

The translation of ATF4 is promoted by eIF2α Ser51 phosphorylation, a hallmark of ISR [62, 63] and by mTORC1, which promotes the translation of ATF4 by mechanisms independent of eIF2α phosphorylation [26, 64]. ISR is induced by ER stress, which activates the eIF2α kinase PERK and by amino acid deprivation which increases the relative abundance of uncharged tRNAs, activating the general control non-derepressible 2 (GCN2) eIF2α kinase [62]. Importantly, the GCN2/p-eIF2a/ATF4 axis is also activated by MYC [60, 63], which is positively regulated by KDM2B. To determine whether KDM2B also promotes ATF4 translation via ISR or amino acid deprivation, we examined the effects of shKDM2B on phosphorylation of eIF2α at Ser51 and we observed a significant reduction (**Fig. 5D**). However, treatment with tunicamycin, which induces ER stress and activates the ISR [65], rescued the phosphorylation of eIF2a, but failed to rescue the expression of ATF4 (**Fig. 5E**), suggesting that KDM2B regulates ATF4 by mechanisms that supersede the effects of eIF2α phosphorylation.

mTORC1, which also regulates the translation of ATF4, is unlikely to contribute to the regulation of ATF4 by KDM2B, because the KDM2B knockdown did not alter the abundance of the activating phosphorylation of mTOR at Ser2448 and Ser2481 (**Fig. 5F**). The preceding data collectively suggest that KDM2B promotes ATF4 expression primarily by augmenting its transcription and not its translation. This suggestion was confirmed by parallel ribosome profiling studies in control and shKDM2B-transduced MDA-MB-231 cells, which showed that the translational efficiency of the ATF4 mRNA is not downregulated by the knockdown of KDM2B [66].

To determine the epistatic relationship between KDM2B, ATF4 and MYC, we knocked them down individually in MDA-MB-231 cells and we examined the effects of the knockdown on their expression and the expression of their enzyme-encoding target genes. The knockdown of KDM2B downregulated MYC and ATF4, and the knockdown of MYC downregulated ATF4, KDM2B and their enzyme targets at both the RNA and protein level (**Fig. 5A,B,G, and S7A,B)**. Moreover, the knockdown of ATF4 did not affect the expression of either KDM2B or MYC, but like the knockdown of KDM2B and MYC, downregulated the expression of the target enzymes (**Fig. 5G**). We conclude that the epistatic order is KDM2B-MYC, ATF4, enzyme targets, with KDM2B and MYC functioning as positive regulators of each other.

Our earlier studies had shown that KDM2B regulates the expression of EZH2, the catalytic component of the PRC2 complex, and that the combined action of KDM2B and EZH2 results in transcriptional repression [8, 10]. Here we showed that, in agreement with earlier studies [7, 9, 15, 45], KDM2B also functions as a transcriptional activator of target genes. To determine whether KDM2B-dependent transcriptional activation also requires the coordinate action of KDM2B and EZH2, we knocked down EZH2 and we observed that it does not result in MYC or ATF4 downregulation (**Fig. 5H**). Therefore, KDM2B does not require EZH2 for the activation of gene expression.

KDM2B is the DNA targeting component of the ncPRC1.1 complex, composed of PCGF1, RING1A/B, BCOR and its homolog BCORL1, SKP1, USP7, RYBP and TRIM27, and linked to transcriptional activation [7, 45]. The knockdown of KDM2B results in the downregulation of not only KDM2B but also the ncPRC1.1 complex members, USP7, TRIM27, SKP1 and RYBP (**Fig. 5I**). We therefore examined whether KDM2B promotes gene expression by functioning in the context of ncPRC1.1. To address this question, we knocked down RING1B, BCOR and PCGF1, and one of the components of the canonical PRC1 complex (PCGF4/BMI1) and we showed that whereas ncPRC1.1 regulates the expression of KDM2B target genes, PRC1 does not (**Fig. 5J, K**).

### KDM2B contributes to the transcriptional activation of enzymes in the SGOC and glutamate metabolism pathways in concert with MYC and ATF4

The preceding data provided evidence that KDM2B, promotes the transcriptional activation of MYC, ATF4, and their downstream metabolic targets, via ncPRC1.1. Given that KDM2B is the DNA targeting component of ncPRC1.1, these data suggested that KDM2B promotes transcription by binding to and regulating the chromatin structure of target genes. Earlier studies had indeed shown that KDM2B-targeted ncPRC1.1 promotes the acetylation of histone H3 at K27 (H3K27ac), a mark associated with transcriptional activation, and the ubiquitination of histone H2A at K119 (H2AK119ub) [45]. To address this question in an unbiased way, we examined the chromatin accessibility of KDM2B target genes in shKDM2B and control MDA-MB-231 cells using ATAC-Seq.

Analysis of the ATAC-Seq data revealed that whereas there were no significant differences in the peak distribution between the control and the KDM2B knockdown cells **(Fig. S8A)**, there were significant quantitative differences, with decreasing accessibility in 25312 sites (Log2FC>=+0.58, FDR<0.05) and increasing accessibility in 12037 sites (Log2FC<=-0.58, FDR<0.05) (**Fig. 6A, S8B)**. Importantly, increasing accessibility in a region up to 2 kb upstream and downstream of the TSS, exhibited an excellent correlation with the upregulation of gene expression. However, the correlation between decreasing chromatin accessibility and decreasing expression was relatively weak (**Fig. 6B**). This could potentially be explained by the dependence of the expression of these genes on diffusely distributed chromatin accessible sites, that are not limited to the TSS, and include enhancers and super enhancers (SEs). The changing accessibility in SEs was confirmed with the analysis of SE associated peaks, which identified 1365 peaks with decreasing accessibility and 654 peaks with increasing accessibility in shKDM2B cells. The importance of diffusely distributed chromatin accessibility changes is illustrated by our data on peak distribution in MYC, ATF4 and genes encoding KDM2B/MYC/ATF4 enzyme targets. These data indeed show that, in shKDM2B cells, accessibility is decreased in the MYC promoter and SE and in the ATF4 enhancer. In addition, reduced accessibility was observed within or upstream the genes encoding PSAT1, MTHFD1L, PHGDH, MTHFD2 and in the SE of MTHFD1L (**Fig. 6C,D**). Alternatively, small drops in chromatin accessibility in promoters that remain accessible, may not be sufficient to prevent the binding and activity of transcription factors targeting these genes.

**Figure 6.**
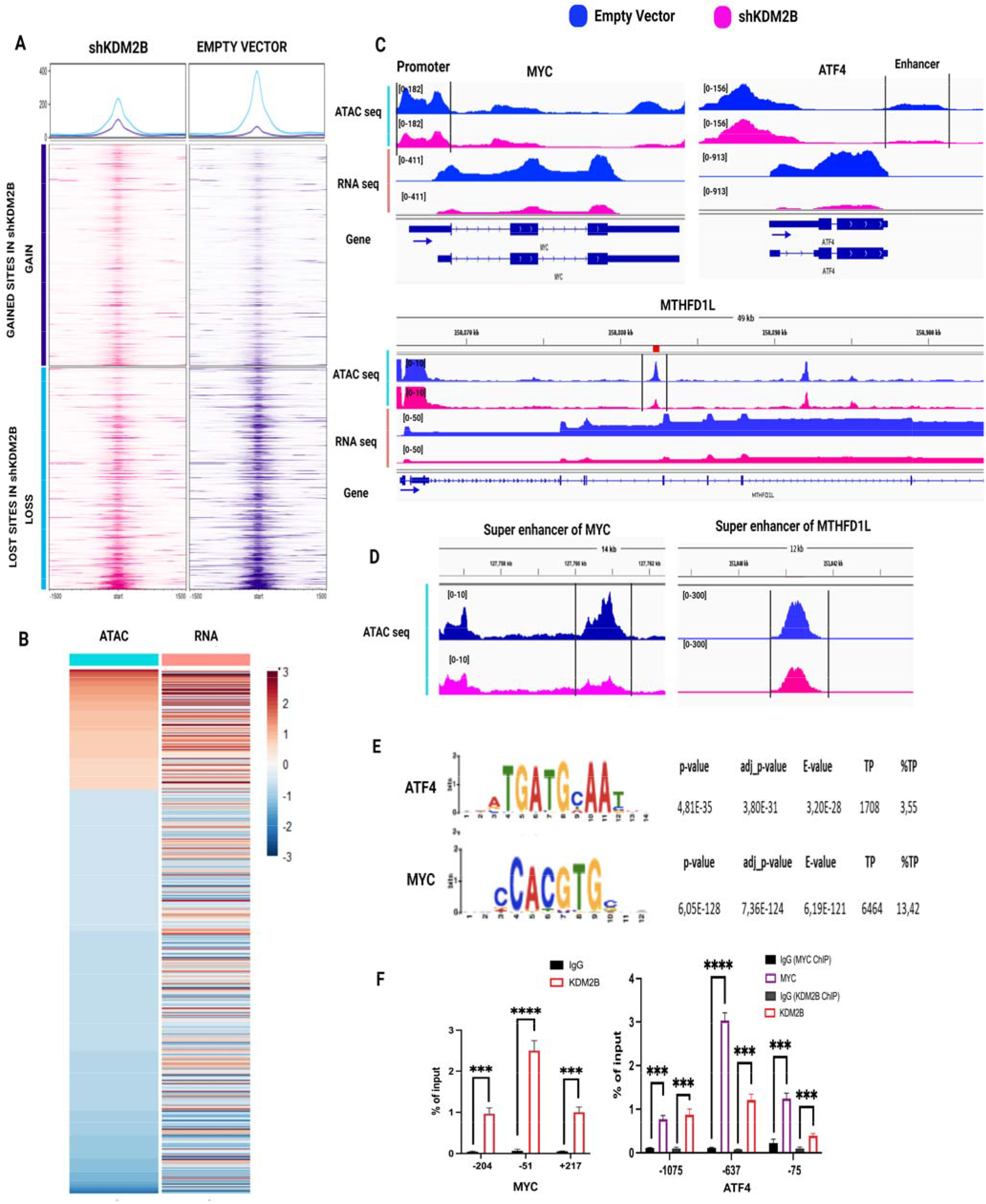
The knockdown of KDM2B alters chromatin accessibility of multiple genes, including MYC and ATF4, and binds the MYC promoter and the ATF4 promoter, in concert with MYC. **A)** Heatmaps and density plots of all regions with increasing or decreasing accessibility in shKDM2B-transduced MDA-MB-231 cells. (Restricted to ±1.5-kb around the TSS). **B)** Heatmaps showing shKDM2B-induced changes in the expression of genes whose promoters (±2-kb around the TSS) exhibit increasing or decreasing chromatin accessibility in shKDM2B cells. **C)** ATAC-seq and RNA-seq tracks of the MYC, ATF4 genes and MTHFD1L genes in Empty Vector and shKDM2B-transduced MDA-MB-231 cells. The accessibility of the MYC promoter (top-left panel), ATF4 enhancer (GeneHancer identifier, <u>GH22J039518</u>,) (top-right panel) and the body of the MTHFD1L gene (bottom panel) was reduced in shKDM2B cells. **D)** ATAC-seq tracks showing that chromatin accessibility in the SEs of MYC (peak at chr8: 127760718-127761118, SE id: SE_02_064000060, SEdb database) (left panel), and MTHFD1L (peak at chr6: 151040977-151041377, SE id: SE_02_048100611, SEdb database) (right panel) is decreased in shKDM2B cells. **E)** ATF4 and MYC motifs are among the transcription factor motifs that are significantly enriched in sites of decreasing chromatin accessibility in shKDM2B cells. Enrichment was determined with the “Analysis for Motif Enrichment” (AME) tool. **F)** ChIP-qPCR analyses showing the binding of KDM2B to the MYC promoter (left panel) and the binding of KDM2B and MYC to the ATF4 promoter (right panel). Data show the DNA immunoprecipitated with IgG (control), or the test antibody (MYC or KDM2B) as a percentage of the input DNA. Mean ± SD of three replicates (n=3). Statistical significance was determined with the unpaired two-tailed t-test.

Following the analyses linking chromatin accessibility to gene expression, we used Analysis of Motif Enrichment (AME) to identify transcription factor binding motifs that were either positively or negatively enriched in genes undergoing changes in chromatin accessibility in shKDM2B-transduced cells. The results showed that the binding motifs for MYC and ATF4 were highly enriched in sites of decreasing accessibility (**Fig. 6E**).

The functional relationship between KDM2B and the epistatically ordered MYC and ATF4 genes may be linear, each of them regulating its downstream target. ChIP experiments support this hypothesis by showing that KDM2B binds the MYC and ATF4 promoters, the latter together with MYC (**Fig. 6F**). Alternatively, it may be more complex, with KDM2B, an epigenetic regulator that functions in the context of ncPRC1.1, also controlling the transcriptional output of MYC and ATF4. The latter hypothesis is supported by the preceding data, showing that MYC and ATF4 motifs are significantly enriched in sites with decreasing accessibility in KDM2B knockdown cells and by the ChIP data in Figure 6F demonstrating that KDM2B and MYC bind the promoter of ATF4 in concert. To address the potential contribution of KDM2B to the regulation of MYC and ATF4 target genes in human breast cancer, we employed a bioinformatics strategy to examine their expression in the FUSCC cohort [46, 47]. To this end, we stratified TNBCs into clusters that express high or low levels of KDM2B. Following this, we identified tumors that express similar levels of MYC and ATF4 in the high and low KDM2B groups and we performed differential gene expression analysis comparing the two groups. The results revealed that the expression of 13.3% of the MYC target genes is KDM2B dependent **(Fig. S8C)**.

### Co-occupancy of the promoters of transcriptionally active genes by KDM2B, MYC and ATF4

The preceding data raised the question whether KDM2B MYC and ATF4 bind regulatory DNA elements in their target genes in concert. To address this question, we carried out ChIP-seq experiments, focusing on the distribution of KDM2B, MYC and ATF4 in parental MDA-MB-231 cells. KDM2B functions in the context of ncPRC1.1 and ncPRC1.1 targets are strongly enriched for H3K4me3, and to a lesser extent H2AK119ub and they are devoid of H3K27me3 [7]. Therefore, we also examined the distribution of these marks along with H3K27ac, the histone mark of active promoters **(Fig. S9A,B)**. Analysis of the data identified three distinct gene clusters, with cluster 2 characterized by the concerted binding of KDM2B, MYC and ATF4 in the vicinity of the TSS. The same cluster was also characterized by the high abundance of H3K4me3 and H3K27ac, which are associated with actively transcribed genes, and by the low abundance of H3K27me3, which is associated with transcriptionally inactive genes. The other two clusters were characterized by the low binding of KDM2B, MYC and ATF4 near the TSS, and by the high abundance of histone marks associated with transcriptional repression (H3K27me3 and H2AK119ub, cluster 3) or by the low abundance of all histone marks (cluster 1) (**Fig. 7A**). Venn diagram of the genes, in cluster 2, with KDM2B, MYC, ATF4 and H3K27ac peaks (TSS ± 3kb), confirmed that the majority of these genes are simultaneously targeted by KDM2B, MYC and ATF4 and they carry H3K27ac marks. Importantly, many of the genes encoding enzymes of the SGOC and glutamate metabolic pathway belong to this group (**Fig. 7B**).

**Figure 7.**
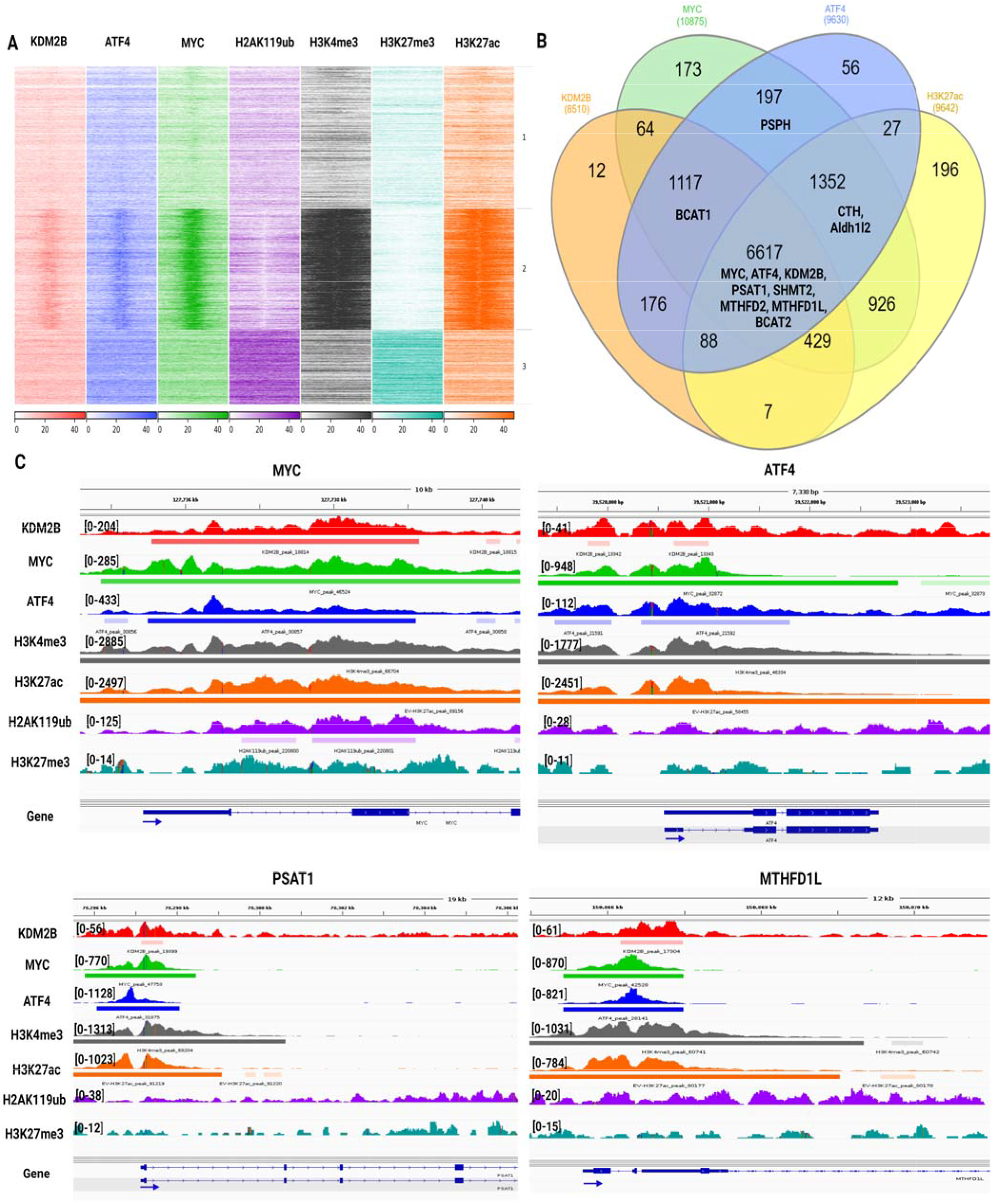
KDM2B, MYC and ATF4 bind in concert the promoters of transcriptionally active genes. **A)** Supervised peak clustering analysis, using the FLUFF analysis tool, and based on promoter occupancy by KDM2B, MYC, and ATF4 (using the ChIP-seq reads-bam files). The analysis identified three clearly defined clusters. Cluster 2 is characterized by extensive overlaps between KDM2B, MYC, and ATF4 binding. Genes in the same cluster are also enriched in H3K4me3 and H3K37ac and they are devoid of H3K27me3 and H2AK119ub marks. Cluster 1 is characterized by low binding of KDM2B, MYC, and ATF4 and by the low abundance of all histone marks, and cluster 3 is characterized by low binding of KDM2B, MYC, ATF4, and high abundance of the repressive marks H2AK119ub and H3K27me3. KDM2B, MYC, ATF4, H3K4me3, H2AK119ub, and H3K27me3 ChIP-seq was performed using two replicates of MDA-MB-231 parental cells. H3K27ac ChIP-seq was performed using two replicates of MDA-MB-231 Empty vector cells. **B)** Venn diagram displaying the frequency of cluster 2 genes whose promoters display different combinations of KDM2B, MYC, or ATF4 binding and H3K27ac abundance (±3 kb around the TSS). **C)** Representative ChIP-seq tracks showing the binding of KDM2B, MYC and ATF4 and the abundance of histone H3K4me3, H3K27ac, H2Ak119ub, and H3K27me3 marks in the promoter region and the gene body of MYC, ATF4 and the SGOC genes PSAT1 and MTHFD1L.

Focusing on the ChIP-seq data in the MYC and ATF4 promoters confirmed that KDM2B also binds both promoters in concert with MYC and ATF4 (**Fig. 7C**), in agreement with the ChIP-qPCR results in figure 6F. Furthermore, in agreement with the data in figures 7A and 7B, H3K4me3 and H3K27ac were highly abundant in both promoters. Representative ChIP-seq data focusing on two of the SGOC targets (PSAT1 and MTHFD1L) showed similar results (**Fig. 7C**).

### Basal-like TNBCs with high expression of KDM2B, MYC and ATF4 exhibit a distinct metabolic signature which is known to carry poor prognosis

Earlier studies by Gong and colleagues [47] on the FUSCC TNBC Cohort, revealed that the 360 primary tumors with RNA-Seq data could be clustered into three metabolic pathway-based subtypes, MPS1, MPS2 and MPS3. The clustering was based on the activity of 86 metabolic pathways, which was calculated from the relative expression of 1660 human genes assigned to these pathways in the KEGG database. Of these clusters, MPS1 was characterized by upregulation of lipid biosynthesis, MPS2 by upregulation of carbohydrate, nucleotide and amino acid metabolism and MPS3, also referred to as the mixed subtype, was characterized by upregulation of combinations of these pathways. Importantly, the MPS2 cluster was characterized by higher tumor grade and poor prognosis relative to the other two clusters. To determine whether KDM2B is associated selectively with any of these subtypes, we analyzed the available transcriptomic data and we observed that although its expression exhibits significant variability in all subtypes, it is significantly higher in basal-like MPS2 and MPS3. MYC expression was also higher in the same subtypes (highest in MPS2) and ATF4 was higher only in MPS2 (**Fig. 8A**). Additionally, the expression of the ncPRC1.1 members, PCGF1 and USP7, was also highest in the MPS2 tumors (**Fig. 8B**). In agreement with data presented in the preceding sections, these results suggest that KDM2B selectively contributes to the MPS2 phenotype, in concert with MYC and ATF4, in the context of ncPRC1.1. To test this hypothesis, we examined the expression of KDM2B, MYC and ATF4 or KDM2B, MYC, ATF4, PCGF1 and USP7, across all basal-like tumors in the three metabolic subtypes and we observed that all these genes tend to be expressed at higher levels in the MPS2 subtype in concert (**Fig. 8C,D**). Importantly, tumors in this metabolic cluster also express the highest levels of a set of KDM2B/MYC/ATF4 metabolic targets (**Fig. 8E**).

**Figure 8.**
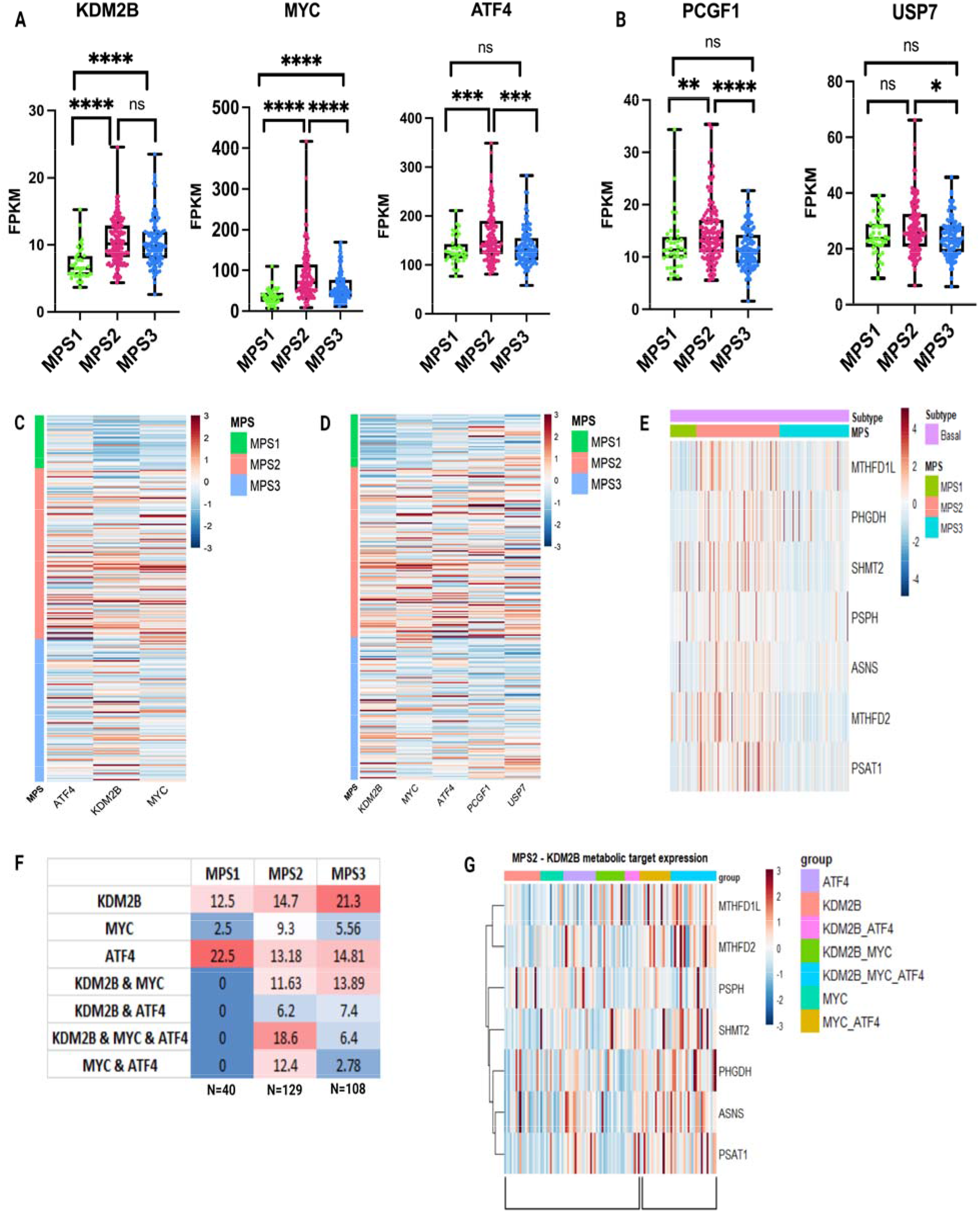
Basal-like TNBCs expressing high levels of KDM2B, MYC and ATF4 exhibit a distinct, prognostically unfavorable metabolic signature. **A)** Box plots of the expression of KDM2B, MYC and ATF4 in the MPS1, MPS2 and MPS3 metabolic subtypes of basal-like TNBCs of the FUSCC cohort. The expression of all three factors tends to be higher in tumors of the MPS2 subtype. Statistical analyses were performed using the Wilcoxon Signed Rank Test. **B)** Box plots of the expression of the ncPRC1.1 complex members, PCGF1 and USP7, in the MPS1, MPS2 and MPS3 metabolic subtypes of basal-like TNBCs of the FUSCC cohort. The expression of both ncPRC1.1 components tends to be higher again in tumors of the MPS2 metabolic subtype. Statistical analyses were performed using the Wilcoxon Signed Rank Test. **C)** Heatmap comparing the expression of ATF4, KDM2B and MYC in the MPS1, MPS2 and MPS3 metabolic subtypes in basal-like TNBCs, in the FUSCC cohort. Concerted upregulation of all three factors in the MPS2 tumors. **D)** Heatmap comparing the expression of KDM2B, MYC, ATF4 and the ncPRC1.1 components, PCGF1 and USP7 in the MPS1, MPS2 and MPS3 metabolic subtypes in basal-like TNBCs, in the FUSCC cohort. Concerted upregulation of all factors in the MPS2 tumors. **E)** Heatmap comparing the expression of KDM2B-regulated genes encoding enzymes of the SGOC and glutamate metabolism pathways, in the MPS1, MPS2 and MPS3 metabolic subtypes in basal-like TNBCs, in the FUSCC cohort. Concerted upregulation of all these KDM2B/MYC/ATF4-regulated genes in MPS2 tumors. **F)** Percentage of tumors (%) in the MPS1, MPS2 and MPS3 metabolic subtypes expressing high levels of KDM2B, MYC and ATF4 in different combinations. High expression was defined as expression (FPKM) equal to or higher than the median. N: Number of tumors in each MPS group. **G)** Heatmap comparing the expression of genes encoding MPS2-associated enzymes in different KDM2B/MYC/ATF4 tumor groups.

Data in this report suggest that the high expression of genes encoding MPS2-associated metabolic enzymes depends on the concerted overexpression of KDM2B, MYC and ATF4. We therefore hypothesized that the MPS2 metabolic subtype will contain the highest percentage of tumors with concerted overexpression of all three factors. To address this hypothesis, we divided the tumors in the three metabolic subtypes into groups expressing high levels of KDM2B, MYC and ATF4 in all possible combinations and we observed that the highest percentage of triple positive tumors indeed belonged to the MPS2 subtype (18.6%) (**Fig. 8F**). High expression was defined as expression equal to, or higher than the median expression. MPS2 also contained the highest percentage of tumors with high expression of MYC and ATF4 (12.4%) (**Fig. 8F**). However, 8/16 tumors in this subgroup expressed KDM2B at levels slightly below the median and could therefore be functionally equivalent to the triple positive tumors. To determine whether the expression of the MPS2-associated SGOC enzymes depends on the concerted activities of KDM2B, MYC and ATF4, we examined the expression of the genes encoding these enzymes in tumors that belong to different KDM2B/MYC/ATF4 groups and we observed that triple positive tumors indeed expressed the highest levels (**Fig. 8G**). MYC/ATF4 high tumors also express high levels of these enzymes, although as mentioned earlier, half of these tumors may also be functionally triple positive.

## Discussion

Data in this report confirmed that KDM2B protects not only normal [15], but also cancer cells from oxidative stress by activating antioxidant mechanisms. Specifically, KDM2B regulates the abundance of GSH and NADPH, and its knockdown sensitizes cells to ROS inducers, GSH targeting molecules and DUB inhibitors, as expected. However, our attempts to link the regulation of the abundance of GSH by KDM2B to the immediate regulators of GSH synthesis and utilization, were unsuccessful. To address this problem, we resorted to non-biased transcriptomic, quantitative proteomic and metabolomic analyses. These analyses not only addressed this question, but also significantly expanded our understanding of the role of KDM2B in the regulation of intermediary metabolism.

Although KDM2B exerts global effects on metabolism, this report focuses on its role in the SGOC pathway and the interconnected transsulfuration, glutamate and glutathione biosynthesis pathways, which are known to play a major role in stem cell self-renewal, cell proliferation, and cancer [21, 24, 38, 39, 50, 67]. These pathways form a network, which is organized around two interconnected methylation cycles that use folate as a carrier of methyl groups and operate in the mitochondria and the cytoplasm (Fig. 3A). The network is modular, and one of its modules, the synthesis of purines and pyrimidines, is upregulated globally in essentially all cancers [23, 26]. Other modules, including the synthesis of serine and several additional amino acids, as well as the synthesis of GSH and NADPH, exhibit variable activity in different tumors, suggesting that they have context-dependent roles [50]. The role of KDM2B in the regulation of these pathways was demonstrated by the effects of its knockdown on the abundance of the enzymes that control their activities and on the abundance of pathway-associated metabolites (**Fig. 3A**). The results showed that KDM2B regulates primarily the mitochondrial methylation cycle, which is also the cycle primarily deregulated in cancer cells [23, 26, 27, 28, 29]. Its regulation by KDM2B was suggested by our observation that the KDM2B knockdown selectively downregulates the enzymes controlling the mitochondrial cycle, as well as sideroflexins 2 and 3 (SFXN2 and SFNX3, but not SFXN1), which are required for the mitochondrial uptake of serine [68], and the solute carrier SLC25A38, which is required for the mitochondrial uptake of glycine [28]. Based on these observations we conclude that KDM2B regulates the cancer relevant mitochondrial cycle by targeting both the uptake of the molecules feeding this cycle and the abundance of the enzymes that control the utilization of these molecules.

The inhibition of serine biosynthesis, a central component of the metabolic network explored in this report, can be explained from the downregulation of PHGDH, PSAT1 and PSPH in the KDM2B knockdown cells. However, the pathway may also be inhibited because of defects in glucose utilization. The increase in the abundance of Sorbitol-6-phosphate and N-Acetyl-galactosamine is consistent with a shift in carbohydrate metabolism toward the sorbitol and hexosamine pathways. Moreover, the downregulation of serine may introduce a block in the final stages of glycolysis, as serine allosterically activates PKM2 [69]. Reduction of the activity of PKM2 in low serine cells, will delay the transition of phosphor-enol-pyruvate to pyruvate and will reduce the glycolysis-dependent supply of the TCA cycle, and oxidative phosphorylation.

Data in this report show that the biosynthesis of serine, the major methyl donor in the SGOC pathway, is significantly impaired. The serine biosynthesis defect may be compensated by the uptake of serine from the cellular microenvironment. However, the effects of the KDM2B knockdown on the SGOC/Glutamate metabolic network were observed even in cells growing in media supplemented with high amounts of nonessential amino acids, including serine. This could be explained by earlier observations suggesting that the functional output of the pathway cannot be rescued by exogenous serine, and that PHGDH is required for nucleotide synthesis and tumor growth, even under conditions of high availability of exogenous serine. This has been attributed to a compensatory increase in the activity of SHMT1 in PHGDH low cells, which shifts 1C units to the cytosolic cycle making them unavailable for nucleotide synthesis [57]. Alternatively, the loss of PHGDH affects the balance between the PPP and the biosynthetic activities of the TCA cycle, which support nucleotide synthesis [70]. The link connecting PHGDH with the TCA cycle may be PSAT1, which utilizes 3-phospho hydroxy pyruvate and L-glutamate to generate α-Ketoglutarate [70]. In addition to feeding the TCA cycle, α-Ketoglutarate, and its derivative 2-OH-glutarate control the activities of JmjC domain protein and DNA demethylases, which regulate the epigenome [71]. Exogenous serine may not rescue the functional output of the pathway, also because of defects in serine uptake. However, the role of serine uptake defects in shKDM2B cells is uncertain because shKDM2B promotes competing serine uptake regulatory mechanisms. The downregulation of ASNS and the decrease in asparagine in these cells is expected to inhibit serine uptake [72], while the upregulation of the Na^+^-dependent Ala/Ser/Cys/Thr transporter, SLC1A4, promotes serine uptake [73].

In addition to serine, which is the major donor of 1C carbon units, such units can also be donated by glycine, tryptophan, and histidine [23]. However, the ability of any of these amino acids to replace serine as a donor of 1C units in MDA-MB-231 cells is unlikely. Although glycine donates 1C units to THF via decarboxylation catalyzed by the GLDC, MDA-MB-231 cells do not express GLDC. We should add that even if glycine could be utilized as a donor, it would not be expected to rescue the proliferation defect in serine deprived cells [21]. Similarly, some of the enzymes involved in the utilization of tryptophan and histidine are not expressed or they are expressed at very low levels in these cells making their potential contribution to the supply of one carbon units unlikely.

The shKDM2B-induced downregulation of the enzymes involved in formate synthesis suggests that KDM2B knockdown cells are formate deficient and this was confirmed by direct measurement of formate levels. In addition to the formate deficiency, these cells are also defective in PPP, as evidenced by the low levels of Glyceraldehyde-3-phosphate, Erythrose-4-phosphate, and Ribose-5-phosphate-isomerase A (RPIA) an enzyme that catalyzes the synthesis of Ribose 5-phosphate. Formate and other 1C carbon units and ribose-5-phosphate are the building blocks for nucleotide synthesis [23, 24]. Their decrease should therefore impair nucleotide and nucleic acid production and is likely to be the main factor responsible for the inhibition of cellular proliferation by shKDM2B.

Data presented in figure 4E show that KDM2B knockdown cells exhibit a significant reduction in OCR. The defect in mitochondrial oxidative phosphorylation, which is suggested by these data can be caused by metabolic shifts that impair the translation of mitochondrial mRNAs [58], as well as by defects in sphingolipid metabolism [74], both of which are deregulated in KDM2B knockdown cells. Specifically, shKDM2B downregulates SHMT2, MTHFD2 and MTFMT, which promote the formylation of the initiating methionine tRNA, that is required for translational initiation of mitochondrial proteins involved in the electron transport chain [58]. Mitochondrial mRNA translation defects may also be caused by the impaired utilization of 5,10-Methylene-THF, which contributes to the generation of the taurinomethyluridine base of the lysine and leucine tRNAs [75].

The contribution of KDM2B in the biosynthesis of ceramides was suggested by observations in this report showing that the knockdown of KDM2B decreases the availability of serine and downregulates the expression of SPTLC3, a member of the serine-palmitoyl-transferase complex, which catalyzes the first rate limiting step in the de novo synthesis of sphingoid bases and ceramides [76]. The mitochondrial defect in the shKDM2B-transduced, SGOC-deficient cells, combined with the previously observed increased sensitivity of serine-deprived cells to mitochondrial complex I targeting biguanides [77] suggest that KDM2B knockdown cells may be sensitive to metformin.

Data presented in this report provide insights into the molecular mechanisms responsible for several aspects of the KDM2B phenotype, including the neural tube closure defect of the KDM2B knockout mice [40], the promotion of stem cell self-renewal [10] and the promotion of angiogenesis [8]. The neural tube closure defect is probably due to the mitochondrial dysfunction induced by the loss of KDM2B. The *KDM2B* gene is transcribed from multiple alternative promoters, and different transcripts are translated into distinct protein isoforms [2]. One of these isoforms lacks the N-terminal JmjC domain (short isoform). Knocking out both the long and the JmjC domain-deficient short isoform results in embryonic lethality in mice, while knocking out only the JmjC domain sufficient long isoforms results in live mice with neural tube closure defects [40]. Such defects are also induced by folate deficiency and by mutations affecting 1C metabolism [37], both of which impair basal mitochondrial respiration. The neural tube defects of these mice are therefore compatible with the results presented in this report.

Our earlier studies had shown that KDM2B promotes cancer stem cell self-renewal by regulating the expression of multiple members of the polycomb complexes [8, 10] and parallel studies by others have reported that the SGOC pathway is also required for the maintenance of the stem cell phenotype [38, 39]. Finally, our earlier studies had provided evidence that the regulation of angiogenesis by FGF2 depends on KDM2B [8], while work by other groups had shown that the SGOC metabolic pathway also promotes angiogenesis [78]. These observations combined, suggest that the role of KDM2B in angiogenesis may also depend on its role in SGOC regulation. Other cancer relevant phenotypes that are regulated by the SGOC pathway and they may therefore depend on KDM2B include tumor cell invasiveness [79] and inflammation [79, 80].

KDM2B is a histone demethylase that contributes to the regulation of gene expression and our data confirmed that its global effects on metabolism are due to its role in the transcriptional regulation of a host of metabolic enzyme genes. Its role in the regulation of these genes depends on ATF4 and MYC, both of which are transcriptionally regulated by KDM2B. Regarding MYC however, several pieces of evidence suggest that KDM2B may regulate not only its expression, but also its transcriptional activity: a) analysis of ChIP-seq data from MDA-MB-231 cells revealed that KDM2B binds the promoters of many active genes, including those of MYC, ATF4, and many of their targets, together with MYC (and ATF4). These data suggest that there is a set of transcriptionally active genes, whose expression depends on the concerted action of KDM2B, MYC (and ATF4). b) AME analysis of ATAC-Seq data from the same cells revealed significant enrichment of MYC (and ATF4) motifs in KDM2B-dependent chromatin-accessible TSS sites, suggesting that KDM2B may render these sites accessible to MYC (and ATF4) binding; c) The highest expression of KDM2B, MYC and ATF4 among all tumors of the FUSCC cohort was observed in the MPS2 metabolic subtype. The same metabolic subtype also contained the highest percentage of tumors expressing high levels (above the median) of all three factors in concert. Importantly, the tumors in the MPS2 metabolic subtype also expressed high levels of a set of KDM2B, MYC and ATF4 metabolic targets, d) Finally, comparison of the FUSCC cohort tumors expressing high or low levels of KDM2B but similar levels of MYC and ATF4, identified a set of MYC target genes whose expression correlates with the expression of KDM2B, suggesting that their regulation by MYC is KDM2B-dependent.

Although KDM2B functions either as a transcriptional repressor or transcriptional activator, it is its transcriptional activator function that regulates the enzyme-encoding genes in this report. Transcriptional repression by KDM2B is mediated through the concerted action of KDM2B and EZH2, which based on the data in this report has no role in KDM2B-dependent transcriptional activation. Given that KDM2B is also the targeting component of the ncPRC1.1 complex, we questioned whether its transcription activator function is ncPRC1.1-dependent and the results confirmed the hypothesis. The activation of transcription by KDM2B in concert with MYC and ATF4 in the context of ncPRC1.1 suggests that ncPRC1.1 also promotes transcriptional activation via MYC/ATF4. This is compatible with results of earlier studies, which had shown that transcriptional activation of a select group of genes by ncPRC1.1 depends on MYC [7].

Transcriptional regulators of the SGOC enzymes include, in addition to ATF4 and MYC, the Peroxisome proliferator-activated receptor γ coactivator-1α (PPARGC1A, also known as PGC1α) and its downstream target Orphan oestrogen related receptor (ESRRA, also known as ERRα), the Epithelial splicing regulatory protein-1 (ESRP1), the sterol regulatory element binding transcription factor 1 (SREBF1) and the epigenetic regulators G9A (KMT1C/EHMT2) and KDM4C (GASC1) [28]. The role of these proteins in SGOC regulation by KDM2B has not been addressed directly. However, PPARGC1A which is upregulated by shKDM2B, has been shown to inhibit ATF4 expression in oestrogen-positive breast cancer, and the remaining regulators are either not expressed or their expression is not affected by shKDM2B making them unlikely candidates for SGOCP regulation.

In summary, evidence presented in this report linked KDM2B, ATF4 and MYC in a transcriptional network which promotes the expression of genes encoding multiple metabolic enzymes. As a result, KDM2B contributes to the global regulation of intermediary metabolism. The work presented here focuses primarily on the role of KDM2B in the SGOC, and glutamate metabolism pathways and the results provide functional links between metabolism and some of the known phenotypes associated with KDM2B, including tumorigenesis, neural tube closure defect and the stem cell maintenance phenotype. In addition, it suggests KDM2B-dependent metabolic vulnerabilities of cancer cells, some of which, like the sensitivity to DUB inhibitors, ROS inducers and GSH targeting molecules were studied and confirmed, while others will be addressed in future studies.

## Supporting information

Supplementary Tables of the Methods

## Abbreviations

GSH: reduced Glutathione
GSSG: oxidized glutathione
ROS: Reactive Oxygen Species
DUB: deubiquitinase
SGOC: Serine-Glycine-One-Carbon
ATF4: activating transcription factor 4
ncPRC1.1: non-canonical polycomb repressive complex 1.1
TNBCs: Triple Negative Breast Cancers
JmjC: Jumonji C
TMT: Tandem Mass Tag
ATAC-Seq: Assay for Transposase-Accessible Chromatin sequencing
ChIP: chromatin immunoprecipitation
TSS: transcription start site
Aass: aminoadipate-semialdehyde synthase
Nqo1: NAD(P)H quinone dehydrogenase 1
Prdx4: peroxiredoxin 4
Serpinb1b: serpin family B member 1b
PDH: pyruvate dehydrogenase
ATCase: aspartate carbamoyltransferase
1C: One-Carbon
3PG: 3-phosphoglycerate
SSP: Serine Synthesis Pathway
THF: tetrahydrofolate
EZH2: Enhancer Of Zeste 2 Polycomb Repressive Complex 2 Subunit
TBHP: tert-Butyl hydroperoxide
PL: Piperlongumine
NAC: N-Acetyl-Cysteine
ER: endoplasmic reticulum
GLS: glutaminase
GCLC: glutamate-cysteine ligase catalytic
GCLM: glutamate-cysteine ligase modifier
GSS: glutathione synthetase
GSR: glutathione reductase
GPX4: glutathione peroxidase 4
GO: Gene Ontology
GSEA: Gene set enrichment analysis
ASNS: asparagine synthetase
BCAAs: branched chain amino acids
BCAT1/2: branched-chain aminotransferase 1/2
8-OH-Gua: 8-hydroxyguanine
SAM: S-adenosyl-methionine
SAH: S-adenosylhomocysteine
PHGDH: Phosphoglycerate dehydrogenase
3-PHP: 3-phosphohydroxypyruvate
3-PS: 3-phosphoserine
PSAT1: phosphoserine aminotransferase 1
PSPH: phosphoserine phosphatase
SHMT1/2: Serine hydroxymethyltransferase 1/2
MTHFD1/2: methylenetetrahydrofolate dehydrogenase 1/2
MTHFD1L: methylenetetrahydrofolate dehydrogenase 1 like
MTHFR: 5,10-methyleneTHF reductase
MTR: methionine synthase
CBS: cystathionine β-synthase
CTH: cystathionine γ-lyase/cystathionase
ALDH1L1/2: Aldehyde Dehydrogenase 1 Family Member L1/2
OCR: Oxygen Consumption Rate
ISR: integrated stress response
PRC1/2: polycomb repressive complex 1/2
SEs: super-enhancers
KEGG: Kyoto Encyclopedia of Genes and Genomes
TCA: tricarboxylic acid
GLDC: Glycine Decarboxylase Complex
MTFMT: Mitochondrial Methionyl-TRNA Formyl-transferase

## Author contributions

EC and PNT conceived the project, designed experiments, and wrote the manuscript. EC performed most of the experiments and analysed data. VA performed experiments and analysed data. ALF carried out bioinformatic analyses (RNA-seq, ATAC-seq, ChIP-seq, FUSCC cohort). JA performed the metabolomics analysis. GC and MS performed the qRT-PCR and ChIP-qPCR experiments. IC generated the Ridge-line plots and MITHril bar graph. BS, SP, AS assisted with animal and some of the gene expression analyses. GN supervised bioinformatics analyses. LS advised on the design of experiments. MAF supervised metabolomics analysis. PNT supervised the study. All authors approved the final manuscript.

## Acknowledgements

We would like to thank Dr. Yue Gong, Dr. Yi-Zhou Jiang and Dr. Zhi-Ming Shao for providing information on the FUSCC TNBC cohort and the helpful discussions.

## Methods

### Cell culture

MDA-MB-231, MDA-MB-468 and Lenti-X 293T cells were grown in Dulbecco’s Modified Eagle’s Medium (DMEM) (Millipore-Sigma, Cat. D5796) supplemented with penicillin/streptomycin (Corning, Cat. 30-002-CI), nonessential amino acids (Corning, Cat. 25-025-CI), l-glutamine (Corning, Cat. 25-005-CI), sodium pyruvate (Millipore-Sigma, Cat. S8636), plasmocin 2.5LJng/μL (Invivogen, Cat. ant-mpp), and 10% fetal bovine serum (Fisher Scientific, Cat. MT35010CV). Cell lines were periodically checked for mycoplasma, using the PCR mycoplasma detection kit (ABM, Cat. G238).

#### Infection with Lentiviral shRNA constructs

shRNAs in the pLKO-puro lentiviral vector, were packaged in Lenti-X 293T Cells (Takara Bio, Cat. 632180) by transient transfection, in combination with the packaging constructs psPax2 (Addgene, Cat. 12260) and pMD2.G (Addgene, Cat. 12259). Transfections were carried out using the Lipofectamine 3000 Transfection Reagent (Thermo Fisher Scientific, Cat. L3000015) and the Opti-MEM Reduced Serum Medium (Fisher Scientific, Cat. 31-985-070), according to the manufacturer’s protocol. The supernatants were collected 48h and 72h after the transfection. MDA-MB-231 and MDA-MB-468 cells were infected with the viral supernatants, in the presence of 8 μg/mL polybrene (Millipore-Sigma, Cat. 107689). Infected cells were selected with puromycin for 48h (Gibco, Cat. A11138) (10 μg/mL). The shRNAs used in the present study are listed in Supplementary Table 10.

#### Serine and glycine deprivatio

MDA-MB-231 and MDA-MB-468 empty vector and shKDM2B cells were cultured in MEM (Gibco, Cat. 21090) supplemented with 10% dialysed FBS (Thermo Fisher Scientific, Cat. A3382001), 1% penicillin–streptomycin (Corning, Cat. 30-002-CI), 5 mM D-glucose (Thermo Fisher Scientific, Cat. A2494001), 65 µM sodium pyruvate (Millipore-Sigma, Cat. S8636), 1X MEM vitamin solution (Thermo Fisher Scientific, Cat. 11120052), 2 mM L-glutamine (Corning, Cat. 25-005-CI), 0.15 mM L-proline (Millipore-Sigma, Cat. P5607), 0.15 mM L-alanine (Millipore-Sigma, Cat. A7469), 0.15 mM L-aspartic acid (Millipore-Sigma, Cat. A7219), 0.15 mM L-glutamic acid (Millipore-Sigma, Cat. G8415) and 0.34LJmM L-asparagine (Millipore-Sigma, Cat. A4159) (-Ser & Gly media). Cells were grown in the media, which is deficient in Serine and Glycine, or in the same media supplemented with 0.4LJmM L-serine (Millipore-Sigma, Cat. S4311) and/or 0.4 mM L-glycine (Millipore-Sigma, Cat. G8790). Cell proliferation was monitored for 4-5 days in the incucyte live-cell imaging and analysis system. All experiments were done in triplicate.

#### Monitoring cell proliferation with the Incucyte live-cell analysis system

MDA-MB-231 and MDA-MB-468 cells were plated in triplicate in 96, 12 or 24 -well tissue culture dishes. Cell proliferation of cells growing under normal culture conditions was monitored every 12/24 h, using the Incucyte S3 Live-Cell Imaging and Analysis System (Essen Biosciences, Ann Arbor, MI). Both cell lines were monitored for 5-6 days.

For the treatments with the ROS inducer PL (MedChemExpress Cat. HY-N2329) and the inhibitors APR-246 (MedChemExpress, Cat. <u>HY-19980</u>) and MI-2 (MedChemExpress, Cat. HY-12276), the cells were plated at 40% confluence and the growth was monitored for 48h.

Images were captured at sequential time points and analyzed using the Incucyte confluence masking software (Essen Biosciences, Ann Arbor, MI), which calculates the surface area occupied by the growing cells, as a percentage of the total surface area of the well. Confluence monitoring was optimized for each cell line to minimize background. To ensure an unbiased analysis, the optimization parameters for a given cell line were also applied to all the derivatives of that cell line. All the inhibitors and the treatments used in the present study are listed in the Supplementary Table 11.

### Reactive Oxygen Species (ROS)

#### Measuring ROS levels

Intracellular ROS levels were measured by flowcytometry, after treatment with the CellROX orange dye, and following the manufacturer’s instructions (Thermo Scientific, Cat. C10493). TBHP (100 μM), was added to the medium 3 hrs prior to the analysis, which was done using the BD FACSCalibur v2.3 Flow Cytometer (BD Biosciences, San Jose, CA). All the experiments were performed in triplicate and the data were analyzed, using the FlowJo v9.3.3 software.

#### Glutathione

Cellular Glutathione was measured using a Fluorimetric Assay Kit (Abcam, Cat. ab205811) and following the manufacturer’s protocol. Briefly, 100-300 thousand cells (counted using a Bio-Rad, TC20 automated cell counter), were resuspended in 100 µL of ice cold 1X mammalian lysis buffer (Abcam, Cat. ab179835) and deproteinized with the deproteinizing sample preparation kit (BioVision, Cat. K808-200). The sample was then incubated with the reaction mix for 15-20 min, and the levels of reduced GSH were measured using a fluorescence microplate reader (Molecular Devices, SpectraMax iD5) at Ex/Em = 490/520 nm. All experiments were performed in triplicate.

#### Protein Oxidation Detection (Oxyblot)

Oxidized proteins were detected using the OxyBlot Protein Oxidation Detection Kit (Sigma, S7150) and following the manufacturer’s protocol. Briefly, 20 μg of protein were derivatize used with DNPH. Following incubation for 15 min at RT the neutralization solution was added and the samples were electrophoresed in polyacrylamide gels and they were transferred to PVDF membranes. The latter were incubated with the primary antibody for 1h at RT, and then with the secondary antibody for 1h at RT. Oxidized proteins were visualized using the Thermo Scientific Pierce ECL 2 Western Blotting Substrate (Thermo Fisher Scientific, Cat. 80196) and scanned, using the LI-COR Fc Odyssey Imaging System (LI-COR Biosciences, Lincoln, NE).

### Metabolomics

Briefly, 50-70% confluent cultures of control and shKDM2B MDA-MB-231 cells (four biological replicates) were first washed rapidly three times with PBS at room temperature. Following this, 1.5-2×10^6^ cells per sample were treated with ice-cold methanol (80% v/v) and they were snap frozen via submergence into liquid Nitrogen for 30 seconds. Subsequently, they were placed on dry ice and allowed to thaw. This step was repeated three times with 10 second vortex-mixing between cycles. At the end, the samples were centrifuged at 11,500 g for 10 min at 4 °C, and the supernatants were collected, lyophilized overnight (∼14 h) and stored at −80 °C.

Dried pellets were resuspended in LC/MS grade water:methanol (95:5) (Optima, Fisher Chemical) with 0.1% formic acid (Thermo Fisher Scientific Cat. 28905). Untargeted metabolomics analyses were performed using a Q-TOF (Agilent, Cat. 6545) instrument, coupled to a high-performance liquid chromatography (HPLC) system (Agilent, Cat. 1290 Infinity LC). Samples (5 µL) were injected into SB-C18 columns (100 x 2.1 mm, 2.7 µm, InfinityLab Poroshell 120), and two mobile phases were used for chromatographic elution: mobile phase A consisting of LC/MS grade water with 0.1 % formic acid and mobile phase B consisting of LC/MS grade methanol with 0.1 % formic acid. Gradient elution was performed at a flow rate of 0.2 mL/min at 40°C and it was started with 5% mobile phase B, reaching 95% in 15 min. It was then maintained at 95% in mobile phase B for 1 more minute at which point elution was complete. Subsequently, the column was re-equilibrated for 8 min using the initial solvent composition. Three biological replicates were analyzed for each condition in both, positive and negative electrospray ionization (ESI) mode. ESI configuration included a mass range from 100 to 1,200 m/z, full scan mode at a scan rate of 2 scans per second, 3000V of capillary, 10 L/min of nebulizer gas flow and 300 °C of gas temperature. MS/MS data were collected in data dependent acquisition (DDA) mode with a scan rate of 5 spectra/sec and dynamic exclusion of 30 seconds for precursor ion selection and fragmentation, using 10 to 30 V. Mass correction throughout the analysis was achieved by continuous pumping and monitoring of the ions m/z 121.0509 (C5H4N4) and 922.0098 (C18H18O6N3P3F24) for ESI+ mode, and m/z 112.9856 (C2F3O2(NH4) and 1033.9881 (C18H18O6N3P3F24) for ESI-mode.

Raw data conversion to mzML format was performed with ProteoWizard (v3.0.2), followed by feature detection, alignment and deconvolution using OpenMS (v2.6.0). Statistical analyses were performed using Metaboanalyst (v5.0). Statistical significance was defined as FDR < 0.05 for t-test hypothesis testing. Metabolite annotation employed three complementary approaches: 1) accurate mass search, matching m/z with compounds in the Human Metabolome database (HMDB), Kyoto Encyclopedia of Genes and Genomes (KEGG) and Chemical Entities of Biological Interest (ChEBI); 2) spectral search, matching MS/MS fragmentation with mass spectral records in the Global Natural Product Social Molecular Networking (GNPS) and MassBank of North America (MoNA); and 3) de novo molecular structure identification, using SIRIUS (v4.8.2) for structure prediction based on MS and MS/MS isotope and fragmentation pattern analysis.

### Next Generation RNA Sequencing (RNASeq)

#### Library Preparation

RNA was extracted from MDA-MB-231 EV and shKDM2B cells (50-70% confluence) using the Total RNA Purification Plus Kit (Norgen Biotek Corp., Cat. 48300) according to the manufacturer’s protocol. RNA purity and integrity were confirmed using the Agilent ScreenTape (Agilent Technologies Cat 5067-5576) before the preparation of the RNA-Seq libraries. Three biological replicates were used for library preparation. Libraries were prepared, using a KAPA RNA HyperPrep kit with RiboErase (Roche Cat KK8560) and they were analyzed by paired-end sequencing (2×150bp), in an Illumina HiSeq Rapid V2 Chip. Sequencing produced approximately 35 million reads per sample.

#### RNA-Seq analysis

Raw sequencing reads in FASTQ format were quality trimmed, and adapters were removed using Trim Galore (https://www.bioinformatics.babraham.ac.uk/projects/trim_galore/). Trimmed reads were then used as input for RNAdetector (La Ferlita et al., 2021) which integrates alignment, read counting, differential expression analysis, and pathway analysis. Trimmed reads were aligned to the human genome (GRCh38) using STAR (Dobin et al., 2013) and then quantified by featureCounts (Liao et al., 2014). Differential expression analysis was ultimately performed using DESeq2 (Love et al. 2014), edgeR (Robinson et al., 2010), and LIMMA (Ritchie et al., 2015). The differential expression results produced by these three tools were then combined and based on the consensus among all three, we calculated a meta-p-value using metaSeqR (Moulos et al., 2015). Genes with a |Log2FC| > 0.58 (|Linear FC| > 1.5) and a meta-adjusted p-value (Benjamini-Hochberg correction) < 0.05 (determined by metaSeqR) were considered differentially expressed. The identified differentially expressed genes were then used as input to perform a neural-network-based topological pathway analysis using MITHrIL (Alaimo et al., 2016). Pathway shifts with a p-value < 0.05 were considered significant. The potential for differentially expressed genes to be regulated transcriptionally by KDM2B was addressed by exon-intron split analysis using the EISA R package (v.1.8.0) (Gaidatzis et al., 2015). Heatmaps of the results were generated in R using pheatmap (https://cran.r-project.org/web/packages/pheatmap/index.html).

#### Ridgeline-plot

Gene set enrichment analysis of Gene Ontology (GO)-Biological Process terms was conducted using the R package cluster profiler. The input included transcripts that were differentially expressed between Empty Vector and shKDM2B cells with a p-value less than 0.05 and a log2FC (log 2-fold-change) higher than 0.58. The Ridge plot shows the frequency of differentially expressed genes with a log2FC value greater than 0.58 within the top 18 enriched pathways. The pathway names are shown in the y-axis and the mean logFC values of the enriched genes in each GO category are shown in the x axis. Figure 2A focuses on the down regulated processes in the shKDM2B cells, while figure S4A presents both the up-regulated and down-regulated processes. The colour key represents a spectrum of lowest (red) to highest (blue) adjusted p-value value of the enriched terms.

#### Gene Set Enrichment Analysis

Gene Set Enrichment Analysis (v4.3.2) (GSEA) (Subramanian et al., 2005) was performed using the reads per million (RPM) table of the 3 empty vector replicates and the 3 shKDM2B replicates. The geometric mean of the 3 empty vector and 3 shKDM2B samples was calculated for each gene locus and geometric means < 1 were filtered out to remove poorly expressed genes as recommended by GSEA. This resulted in 13250 expressed gene loci, which corresponds to 12538 protein-encoding genes that were used for the analysis. Enrichment analysis was performed, using the Gene Ontology (GO) domain “Biological Process”, which includes terms with >15 and <500 genes per term as the GSEA default. Given that there were only 3 samples in each group (total n=6), the permutations were on the gene list as opposed to the phenotype. 1000 permutations were run for each gene set and an enrichment score was calculated and compared to the actual enrichment score of our input ranked list. Statistics were determined based on the probability that the input ranked list had a better absolute enrichment score than the 1000 permutations and adjusted for multiple hypothesis testing taking into account the total number of considered gene sets.

### TMT proteomics

MDA-MB-231 EV and shKDM2B cells at 50-70% confluency (four replicates of each) were washed in PBS twice, prior to lysis with the proteomics lysis buffer (8M Urea, 50mM triethylammonium bicarbonate) supplemented with Halt Protease and Phosphatase Inhibitor Cocktail (Thermo Fisher Scientific, Cat. 78440). The lysates were sonicated for 30 seconds on ice (amplitude 25%) and following this, the amount of protein per sample was quantified using the Pierce BCA Protein Assay Kit (Thermo Fisher Scientific, Cat. 23225). Trypsin digestion, Tandem Mass Tag (TMT) labelling, fractionation, mass spectrometry, and initial analysis was performed by Bioinformatics Solutions Incorporated (Waterloo, ON, Canada, or website). Briefly, protein (100μg) was reduced, alkylated, precipitated, and then digested with trypsin overnight. Digested samples were then cleaned, using a C18 Stagetip and the peptides in each sample were quantified, using Pierce Quantitative Peptide Assays & Standards (Thermo Fisher Scientific, Cat. 23275). 20μg of each sample was used for TMTpro16plex labelling, and following this, they were pooled and fractionated by high-pH reverse phase chromatography, generating 44 fractions, which were finally combined into 11 fractions. Each fraction was injected twice and analyzed by liquid chromatography–mass spectrometry (LC-MS) via MS3 quantification with synchronous precursor selection. This was done, using an UltiMate™ 3000 RSLCnano liquid chromatography system (Thermo Fisher Scientific, Cat. ULTIMATE 3000 RSLCNANO) attached to a Orbitrap Fusion™ Lumos™ Tribrid™ Mass Spectrometer (Thermo Fisher Scientific, Cat. IQLAAEGAAPFADBMBHQ). Searches of peptide spectra were performed by “Bioinformatics Solutions Incorporated”, using the Uniport validated human proteome and Peaks Studio (v11), the company’s inhouse software (Bioinformatics Solutions Incorporated). The searches were performed using a 15-ppm precursor ion tolerance for peptide mapping. A decoy fusion method was employed to determine if peptide-spectrum matchings are true matchings or likely false positives. A cut-off of 1% false-discovery-rate based on this, was used to filter poor peptide matchings. Statistical significance and fold change was calculated by comparing the peptide quantification values in empty vector and shKDM2B transduced samples, via Peaks Studio (v11).

### Quantitative RT-PCR (qRT-PCR)

Total RNA was extracted using the PureLink RNA Kit (Invitrogen, Cat. No 12183018A). cDNA was synthesised from 1.0 μg of total RNA, using oligo-dT priming and the QuantiTect Reverse Transcription Kit (QIAGEN, Cat No. 205310). Gene expression was measured by quantitative RT-PCR, using the iTaq Universal SYBR Green Supermix (Bio-Rad, Cat. 1725121) and the CFX Opus 384 Real-Time PCR System (Bio-Rad, Cat. 12011452). The qpcr primers used in this study are listed in Supplementary Table 12.

### Western blotting

Cells were lysed in RIPA-buffer (Thermo Fisher Scientific, Cat. 89900) supplemented with phosphatase and protease inhibitor cocktail (Thermo Fisher Scientific, Cat. 78440). Samples were electrophoresed in SDS-PAGE under reducing conditions and they were transferred to a PVDF membrane and analyzed by immunoblotting. Horseradish peroxidase-conjugated goat anti-rabbit or anti-mouse IgG (Cell Signaling Technology, Cat. 7074 and 7076 respectively) were used as secondary antibodies. Proteins were visualized using the Thermo Scientific Pierce ECL 2 Western Blotting Substrate (Thermo Fisher Scientific, Cat. 80196) or the SuperSignal West Femto Chemiluminescent Substrate (Thermo Fisher Scientific, Cat. 34096). Western blots were scanned using the LI-COR Fc Odyssey Imaging System (LI-COR Biosciences, Lincoln, NE). The list of the antibodies used in this study are presented in Supplementary Table 13.

### Metabolite detection and quantification

*<u>Serine and Glycine</u>* concentrations were measured using Serine or Glycine Fluorometric Assay Kits (Abcam, Cat. ab241027 and ab211100 respectively) and following the manufacturer’s protocol. Briefly, lysates derived from 1×10^6^ cells, were deproteinized (removal of enzymes) by pretreatment with the Sample Cleanup Mix and loading to the 10 kD spin columns from BioVision (Cat. 1997-25). Following incubation of the deproteinized samples with the reaction mix for 60min, levels of total serine/glycine were measured on a fluorescent microplate reader (Molecular Devices, SpectraMax iD5) at Ex/Em = 535/587. All experiments were performed in triplicates.

*<u>NADPH</u>* was measured using a Colorimetric Assay Kit (Abcam, Cat. ab186031) and following the manufacturer’s protocol. Briefly, lysates derived from 1.5-2×10^6^ cells were incubated with the reaction mix for 2h at room temperature (RT) in the dark, and levels of NADPH were measured on a microplate reader (Molecular Devices, SpectraMax iD5) at 460 nm. All experiments were performed in triplicate.

*<u>Formate</u>* was measured using a Colorimetric Assay Kit (Sigma, Cat. MAK059-1KT) in lysates derived from 1×10^6 cells. Lysates were incubated with the reaction mix for 60 min at 37°C in the dark, and the levels of formate were measured on a microplate reader (Molecular Devices, SpectraMax iD5) at 450 nm. All experiments were performed in triplicates.

*<u>Glutamate</u>* was measured using the Colorimetric Assay Kit (Abcam, Cat. ab83389) in lysates derived from 2×10^6 cells. Lysates were incubated with the reaction mix for 30 min at 37°C in the dark, and the levels of glutamate were measured on a microplate reader (Molecular Devices, SpectraMax iD5) at 450 nm. All experiments were performed in triplicate.

*<u>S-adenosyl-methionine (SAM)</u>* was measured with a kit from Cell Biolabs (Cat MET-5152). Briefly, 20×10^6^ cells were resuspended in 1ml of cold PBS and sonicated for 6.5 min with 30 sec on, 30 sec off pulses. After spinning at 10.000 x g for 15 min at 4°C, 50 μl of the supernatants were incubated with the primary anti-SAM antibody at RT for 1 hour, and following this, with the diluted HRP-conjugated secondary antibody for an additional hour, also at RT. The SAM-antibody complexes were then incubated with the substrate solution for 5 min, and the reaction was stopped with the addition of stop solution. SAM levels were determined by measuring the absorbance at 450 nm on the SpectraMax iD5 microplate reader. All experiments were performed in triplicate.

Summary of the assays used in this study is presented in Supplementary *Table* S14.

### Tumour xenografts

Control and shKDM2B MDA-MB-231 and MDA-MB-468 cells (2LJ×LJ10^6^ and 4LJ×LJ10^6^ cells respectively), were resuspended in PBS in a total volume of 200 µL and implanted subcutaneously into the flanks of 6-week-old female NSG (NOD-SCID-IL2Rgamma) mice (left side for the empty vector and right side for the shKDM2B cells). The mice were monitored every 4 days and the size of the tumors was measured using a digital caliper. The mice were sacrificed 8 weeks post inoculation. The tumors were resected and weighted. Six NSG mice were used for each cell line.

Mice were housed in a sterile and barrier rodent suite within the Biological Research Tower vivarium of the Ohio State university with controlled temperature, humidity, and with a 12-hours day-night light cycle (6:00 a.m. to 6:00 p.m.). All mice were allowed to free access to water and food.

Ethics statement: All mouse experiments were approved by the Institutional Animal Care and Use Committee (IACUC) of the Ohio State University. IACUC protocol number 2018A00000134, PI: Philip N. Tsichlis.

We used ARRIVEreporting guidelines: Percie du Sert N, Hurst V, Ahluwalia A, Alam S, Avey MT, Baker M, Browne WJ, Clark A, Cuthill IC, Dirnagl U, Emerson M, Garner P, Holgate ST, Howells DW, Karp NA, Lazic SE, Lidster K, MacCallum CJ, Macleod M, Pearl EJ, Petersen O, Rawle F, Peynolds P, Rooney K, Sena ES, Silberberg SD, Steckler T and Wurbel H. The ARRIVE Guidelines 2.0: updated guidelines for reporting animal research.

### Oxygen Consumption Rate (OCR)

Control and shKDM2B cells were seeded on Seahorse XF24 tissue culture plates (Agilent Technologies, Cat. 102342100) the day before the assay. OCR was measured on cells growing at a similar confluence, using the Seahorse XF Cell Mito Stress Test Kit (Agilent Technologies, Cat. 103015-100) and following the instructions of the manufacturer. The assay medium (Agilent Technologies, Cat. 103680100) was supplemented with 10mM glucose (Agilent Technologies, Cat. 103577-100), 2 mM l-glutamine (Agilent Technologies, Cat. 103579-100) and 1mM pyruvate (Agilent Technologies, Cat. 103578-100). The measurement of oxygen consumption was performed with 3 min mixing, 2 min recovery, 3 min measuring, in 3 cycles for 3 hours using the Seahorse XFe24 analyzer system (Agilent Technologies). Inhibitors and substrates were used at the following concentrations: oligomycin (1.5 μM), FCCP (1 μM), Rotenone and antimycin A (0.5 μM). The data were normalized to relative cell confluency, which was measured by the incucyte live-cell imaging and analysis system.

### ATAC-Seq

#### Library preparation

Samples were prepared, using the ATAC-Seq kit (Active Motif, Cat 53150). Briefly, aliquots of 10^5^ MDA-MB-231 EV and shKDM2B cells were centrifuged at 1000LJ×g for 5 min at 4LJ°C. Following one wash with ice-cold PBS cells were resuspended in 100 μl ice-cold ATAC lysis Buffer. 50 μl of Tagmentation Master Mix was added to each sample and incubated at 37LJ°C for 30 min in a thermomixer set at 800 rpm. This was followed by DNA extraction and purification with the DNA Purification Binding Buffer. For library preparation, tagmented DNA was amplified, using unique combinations of i7/i5 indexed primers, and on Illumina’s Nextera adapters. The final step in library purification was a clean-up with SPRI beads (Beckman Coulter, Cat. B23317). The quality of the libraries was confirmed using the Bioanalyzer high sensitivity DNA assay (Agilent Technologies). Libraries were prepared from three biological replicates.

#### Sequencing

Libraries underwent paired-end sequencing (2×150bp) on a Illumina NovaSeq-S4-PE 150 Cycle, which generated an average of approximately 200 million reads per sample.

#### ATAC-Seq Analysis

Raw sequencing reads in FASTQ format were quality trimmed, and adapters were removed using Trim Galore (https://www.bioinformatics.babraham.ac.uk/projects/trim_galore/). Trimmed reads were then aligned to the human genome using Bowtie 2 (v.2.4.5) (Langmead and Salzberg, 2012) and, following this, they were converted into BAM format, sorted for coordinates, and indexed using samtools (v.1.6) (Li et al., 2009). Sorted BAM files were used for peak calling by MACS2 (v.2.2.7.1) (Zhang et al., 2008). Only peaks with a p-value < 0.05 were used for further analyses. Significant peaks were annotated based on their location relative to the TSS of protein-coding genes, by CHIPpeakAnno (v.3.30.1) (Zhu et al., 2010), using the genomic coordinates retrieved from ENSEMBL (internal to CHIPpeakAnno). In addition, significant peaks were annotated relative to super-enhancers, using the genomic coordinates retrieved from SEdb (Wang et al., 2023). Differences in chromatin accessibility were assessed by comparing the peaks in the shKDM2B and EV samples, with the DiffBind R package (https://bioconductor.org/packages/release/bioc/html/DiffBind.html). Peak differences with an adjusted p-value (Benjamini-Hochberg correction) < 0.05 were considered significant. Enrichment in transcription factor binding motifs in peak regions of increasing or decreasing accessibility was assessed using AME (McLeay et al., 2010). To do this, we first used the *getfasta* function of bedtools (v.2.30.0) (Quinlan and Hall, 2010) to retrieve peak sequences in FASTA format from the BED files, which contain the genomic coordinates of the differential peaks. These sequences, in combination with the transcription factor binding motifs retrieved from JASPER (Castro-Mondragon et al., 2022), were finally used as input for AME to run the analysis.

### Chromatin immunoprecipitation

Samples for ChIP-seq experiments were prepared using the SimpleChIP Plus Enzymatic chromatin IP kit (Cell Signaling Technology, Cat. 9005) according to the manufacturer’s protocol. 4×10^6^ MDA-MB-231 cells (per IP) were fixed with the crosslinking agent formaldehyde (Sigma-Aldrich, Cat F8775), (1% formaldehyde for 10 min at room temperature). For the KDM2B ChIP dual cross linking was performed, with 1% formaldehyde, as described above, plus ethylene glycol-bis-succinimidylsuccinate (EGS) (Thermo Scientific, Cat 21565), (1.5 mM in ice-cold PBS). Cells were treated with EGS at RT for 20 minutes. Following chromatin digestion with 0.5 µl Micrococcal Nuclease, the nuclei were lysed with sonication (3 rounds of 20 seconds-on 30 seconds-off pulses with an amplitude of 20%). For chromatin IP, the following antibodies were used: KDM2B 1:100 (Millipore, 17-10264), ATF4 1:100 (Cell Signaling Technology, Cat 11815), c-MYC 1:100 (Cell Signaling Technology, Cat 18583), H3K4me3 1:50 (Cell Signaling Technology, Cat 9751), H3K27me3 1:50 (Cell Signaling Technology, Cat 9733), H2AK119ub 1:100 (Cell Signaling Technology, Cat 8240), H3K27ac (Cell Signaling Technology, Cat 8173). 2% input was used as control.

#### ChIP-qPCR

The immunoprecipitated DNA and the DNA isolated from the 2% input chromatin were amplified by quantitative PCR, using the iTaq Universal SYBR Green Supermix (Bio-Rad, Cat. 1725121) and primers specific for the test loci. Amplification and monitoring was carried out, using the CFX Opus 384 Real-Time PCR System (Bio-Rad, Cat 12011452). Data were presented as the percentage of input, which is the percentage of DNA amplified with primers for a given locus from the immunoprecipitated DNA, relative to the DNA amplified with the same primers from the input sample. The list of the ChIP-qpcr primers used in this study is presented in Supplementary Table 15.

#### ChIP-Seq Library Preparation

ChIP-Seq libraries were prepared using the DNA Library Prep Kit for Illumina (Cell Signaling Technology, Cat 56795) and the Multiplex Oligos for Illumina (Dual Index Primers) (Cell Signaling Technology, Cat. 47538). The starting material for Library construction was: 5 ng of immunoprecipitated DNA for KDM2B, 1 ng for c-MYC and 50 ng for H3K4me3, H3K27me3, and H2AK119ub and 50 ng of the input DNA.

#### ChIP-Seq Sequencing

DNA isolated from the immunoprecipitated samples underwent paired-end sequencing (2×150bp on an Illumina NovaSeq-S4-PE 150 Cycle instrument. This yielded an average of approximately 50-70 million reads per sample.

#### ChIP-seq analysis

Raw sequencing reads in FASTQ format were quality trimmed, and adapters were removed using Trim Galore (https://www.bioinformatics.babraham.ac.uk/projects/trim_galore/). Trimmed reads in FASTQ format were then aligned to the human genome using Bowtie 2 (v.2.4.5) (Langmead and Salzberg, 2012) and, following this, they were converted to the BAM format, sorted for coordinates, and indexed using samtools (v.1.6) (Li et al., 2009). Sorted BAM files of the replicates were merged by samtools (v.1.6) (Li et al., 2009) using the function *merge*. Merged BAM files were finally used as input for MACS2 (v.2.2.7.1) (Zhang et al., 2008) in order to perform the peak calling. Only peaks with a q-value < 0.05 were used for further analyses. Significant peaks were annotated based on their location relative to the TSS of coding-protein genes by CHIPpeakAnno (v.3.30.1) (Zhu et al., 2010) using the genomic coordinates retrieved from ENSEMBL (internal to CHIPpeakAnno). The visualization of the peaks around the TSS and other genomic features was performed using the ChIPseeker R package (v.1.32.1) (Yu et al., 2015) while the clustering of TSS DNA peaks (3KB on either side of the TSS), immunoprecipitated with different antibodies, was performed by Fluff (v.3.0.3) (Georgiou and van Heeringen, 2016) using the Euclidean distance.

### Databases used for the downloading of cancer patients’ data

- The data in Figure 1A (KDM2B expression across the breast cancer subtypes and in normal and Genotype-Tissue Expression (GTEx) samples) were downloaded from the Gepia2 database (http://gepia2.cancer-pku.cn), cut-off: Log2FC>0.6 and p-value <0.05.
- The data in Supplementary Figure 5A were downloaded from the xena browser (https://xenabrowser.net/) using the <u>TCGA Breast Cancer (BRCA)</u> database (24 datasets). The clustering of samples into the Basal-like, Her2 enriched, Luminal A, Luminal B was performed based on the phenotypic PAM50 mRNA subtypes (Cancer Genome Atlas Network, 2012).
- The data in Supplementary Figure 5B (KDM2B correlations with the SGOC genes in the TCGA TNBC tumours) were downloaded from cBioPortal (https://www.cbioportal.org/) using RNA-seq data from the “Breast invasive carcinoma, TCGA Firehose Legacy” database. 116 TNBC patients (ER-, PR-, Her2-).

### 2. The expression of KDM2B, MYC ATF4 and members of the ncPRC1.1 complex in the MPS1, MPS2 and MPS3 metabolic subtypes of the FUSCC TNBC cohort

RNA-Seq raw counts of the FUSCC TNBC cohort were scaled using the Fragments Per Kilobase of transcript per Million mapped reads (FPKM) formula to normalize the differences in depth of coverage across the samples. Box plots showing the differential expression of KDM2B, MYC, ATF4, PCGF1, and USP7 across the metabolic subtypes were generated using GraphPad Prism. Statistical significance was calculated with the Wilcoxon test.

Scaled counts were also used to visualize the expression of KDM2B, MYC, ATF4, PCGF1, USP7, and the SGOC enzymes across the metabolic subtypes in a heatmap generated by pheatmap (https://cran.r-project.org/web/packages/pheatmap/index.html). In the latter, genes were clustered using the Pearson correlation as distance.

Correlations between KDM2B and each of the expressed genes were calculated in R using the Spearman correlation. The correlation p-values were also adjusted for multiple test hypotheses using the Benjamini-Hochberg correction in R. KDM2B correlations with an adjusted p-value < 0.05 were considered significant.

### Analysis of MYC targets in the FUSCC TNBC cohort

The effect of KDM2B on the expression of MYC targets was evaluated by comparing tumors with high and low KDM2B expression and comparable levels of MYC and ATF4 expression. Gene expression (in raw counts) was scaled to normalize differences in depth of coverage across tumors, using the RPM formula. Following this, tumors with KDM2B expression in the third or upper quartile and in the first or lower quartile were identified as high and low KDM2B expressors, respectively. An additional sample filter removed tumor samples that had MYC expression in the second quartile, eliminating tumors with more variable MYC expression. Raw counts of the selected samples were then scaled using the RPM formula and genes with a geometric mean RPM < 1 were removed because they were either not expressed or were expressed at very low levels. Afterward, the raw read counts of the retained genes were log2-transformed with the Voom function and then used for D.E. analysis, leveraging the Limma R package (v3.54.2) (Ritchie et al., 2015). Genes with a |Log2FC| > 0.58 (|Linear FC| > 1.5) and an adjusted p-value (Benjamini-Hochberg correction) < 0.05 were considered differentially expressed, suggesting that the regulation of the expression of these genes by MYC is modulated by KDM2B. The list of MYC targets was retrieved from the Hallmark Gene Sets of GSEA (www.gsea-msigdb.org).

### Statistical analyses

Plots and unpaired two-tailed t test analyses were performed using GraphPad Prism version 9.2.0.

All the Figures including the graphical abstract and the schematic diagrams were created with BioRender.com.”

