## Supplementary Figures S1-9 for "Transcriptional regulation of amino acid metabolism by KDM2B, in the context of ncPRC1.1 and in concert with MYC and ATF4"

**Supplementary Figure 1.**

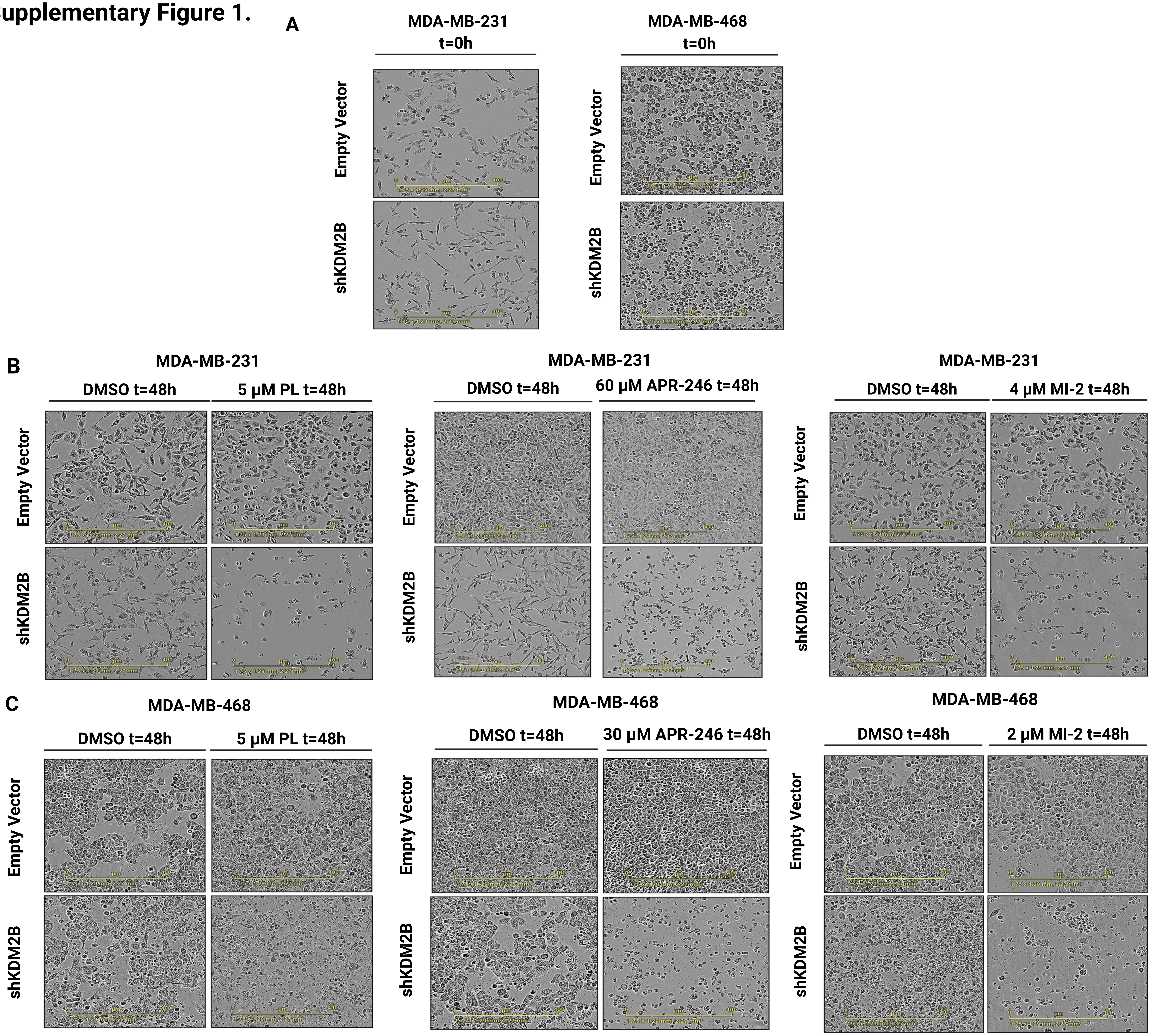


**Supplementary Figure 1. The knockdown of KDM2B inhibits the proliferation of basal-like breast cancer cell lines and enhances their sensitivity to the pro-oxidants PL and APR -246 and the DUB inhibitor MI-2.**

**A-C**. Incucyte-captured images of Empty Vector and shKDM2B-transduced MDA-MB-231 and MDA-MB-468 cells, before (t=0) **(A)** and 48 hours after treatment with DMSO, PL (5 µM), APR-246 (60 µM for MDA-MB-231 and 30 µM for MDA-MB-468 cells), or the DUB inhibitor MI-2 (4 µM for MDA-MB-231 and 2µM for MDA-MB-468 cells) (**B**,**C**).

#### Supplementary Figure 2.

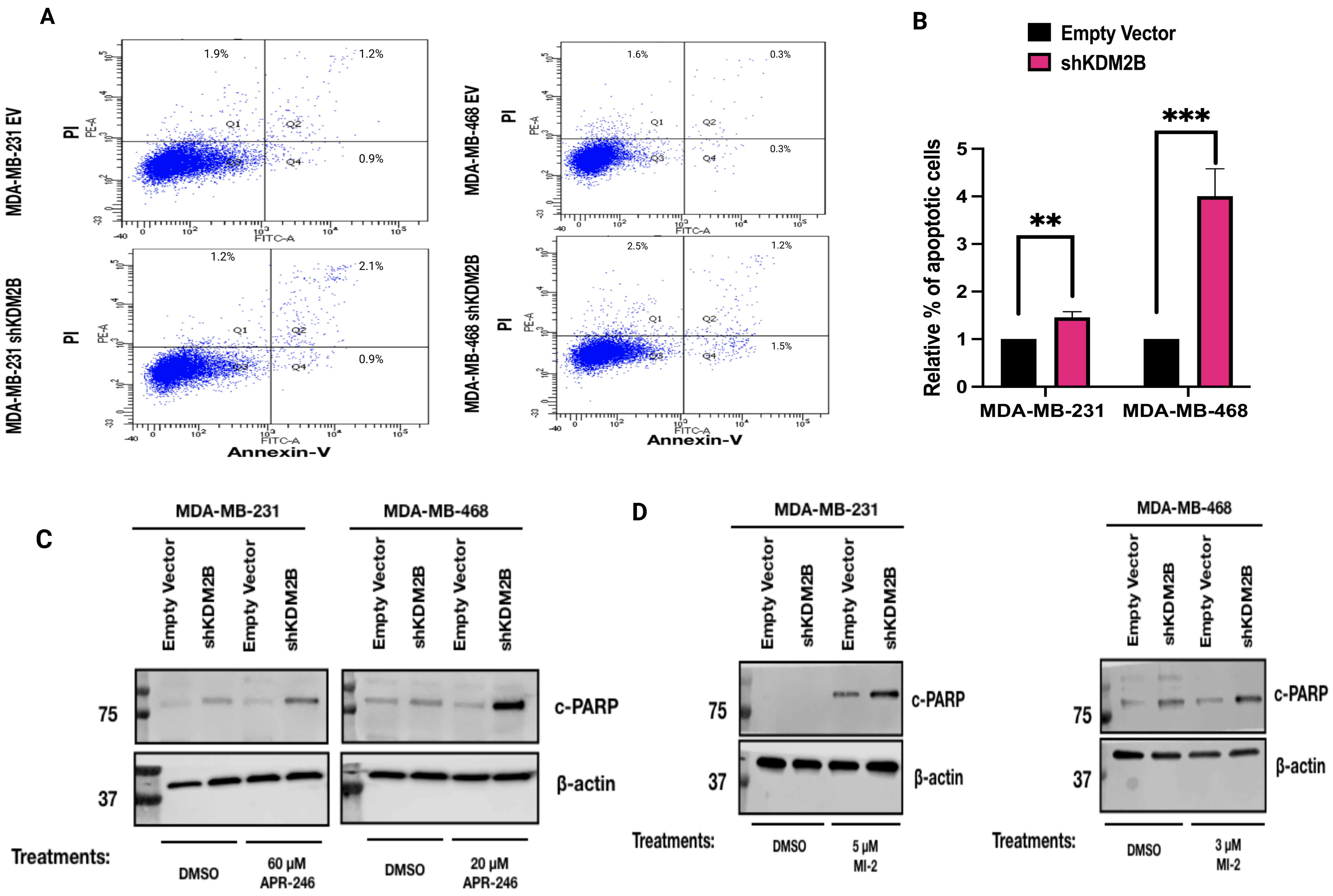


**Supplementary Figure 2. The knockdown of KDM2B promotes apoptosis of basal-like breast cancer cell lines and enhances the sensitivity of such cell lines to the apoptotic activities of APR-246 and MI-2.**
**A)** FACS analysis of Empty Vector and shKDM2B-transduced MDA-MB-231 (left panel) and MDA-MB-468 (right panel) cells stained with Annexin-V and Propidium Iodide (PI). **B)** Quantification of Annexin V positive (apoptotic) cells in both cell lines, transduced with the Empty Vector or shKDM2B (Experiment in A). Mean ± SD of Annexin V-positive cells from one experiment with three biological replicates (n=3). Statistical significance was determined with the unpaired two-tailed t-test. **C)** Immunoblots of Empty Vector and shKDM2B-transduced MDA-MB-231 and MDA-MB-468 cells, treated with DMSO, or APR-246 (60 μM for MDA-MB-231 and 20 μM for MDA-MB-468) for 24 hours, were probed with anti-cleaved-c-PARP antibody. Loading control was β-actin. **D)** Immunoblots of Empty Vector and shKDM2B-transduced MDA-MB-231 and MDA-MB-468 cells, treated with DMSO, or MI-2 (5 μM for MDA-MB-231 and 3 μM for MDA-MB-468 cells), for 24 hours, were probed with the anti-cleaved-c-PARP antibody. Loading control was β-actin.

#### Supplementary Figure 3.

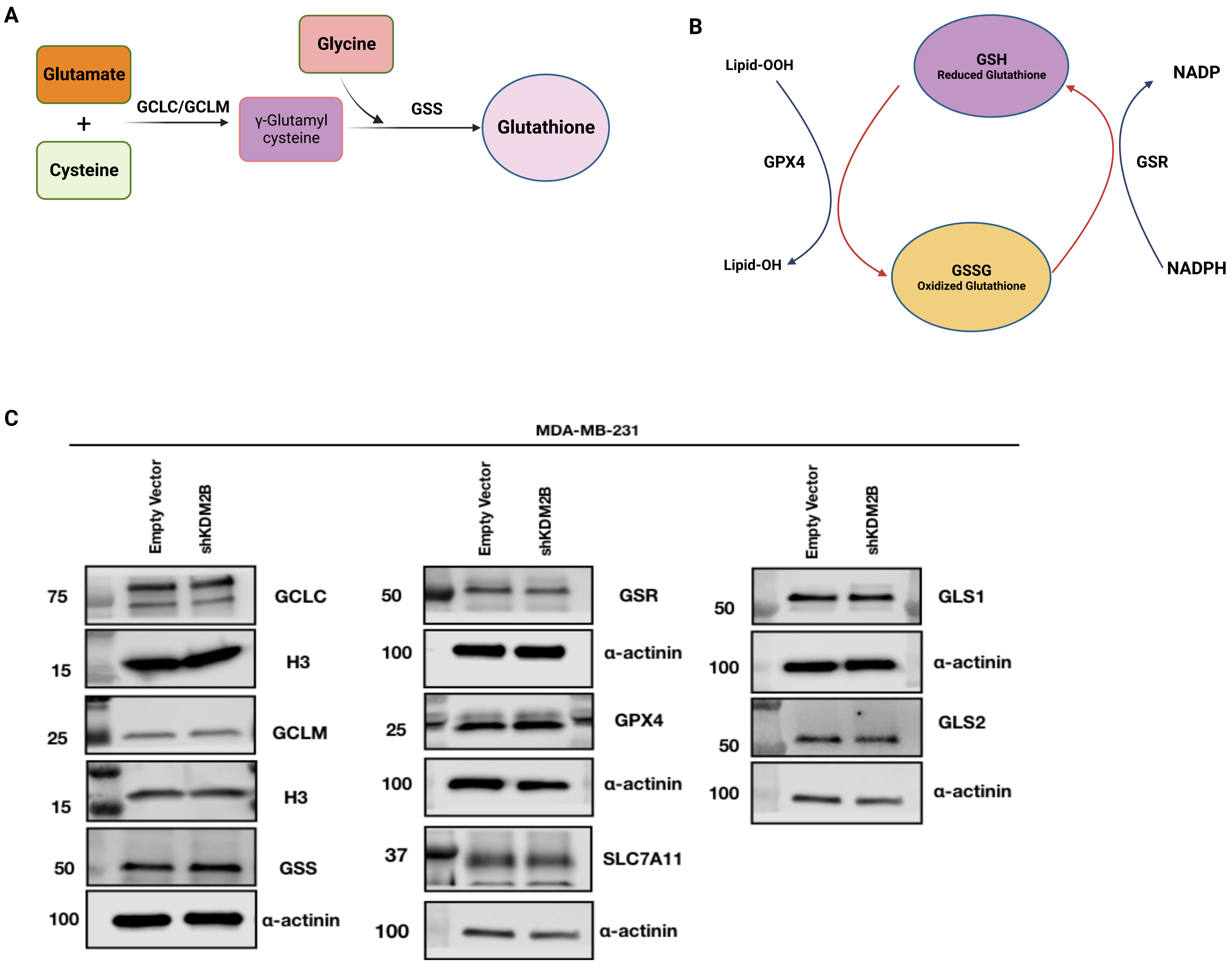


**Supplementary Figure 3. The expression of enzymes and transporters that control the biosynthesis and utilization of GSH, is not regulated by KDM2B.**
**A)** Schematic diagram of Glutathione biosynthesis. Glutathione is a tripeptide, generated from glutamate, cysteine, and glycine in two enzymatic steps. The first is catalyzed by GCL (a complex of GCLC and GCLM), which ligates cysteine to glutamate to produce γ-glutamyl-cysteine. The latter is ligated to glycine via a reaction catalyzed by GSS, to produce Glutathione (GSH). **B)** GSH is the major cellular antioxidant, and it is used by the cells to inhibit or reverse the oxidation of cellular macromolecules, including proteins and lipids. The reduction of ferroptosis-promoting lipid peroxides (Lipid-OOH) to lipid hydroxides (lipid-OH) by GSH, is catalyzed by GPX4. The Oxidized Glutathione (GSSG) is reduced back to GSH by GSR (Glutathione Reductase), via a reaction requiring NADPH. **C)** Immunoblots of Empty Vector and shKDM2B-transduced MDA-MB-231 cells, were probed with antibodies targeting GCLC, GCLM, and GSS (glutathione biosynthesis), GPX4 and GSR (GSH/GSSG interconversion), SLC7A11 (xCt cysteine-glutamate transporter) GLS1 and GLS2, (Glutaminases, promoting the conversion of glutamine to glutamate). Loading controls were α-actinin, or histone H3, as indicated.

**Supplementary Figure 4.**

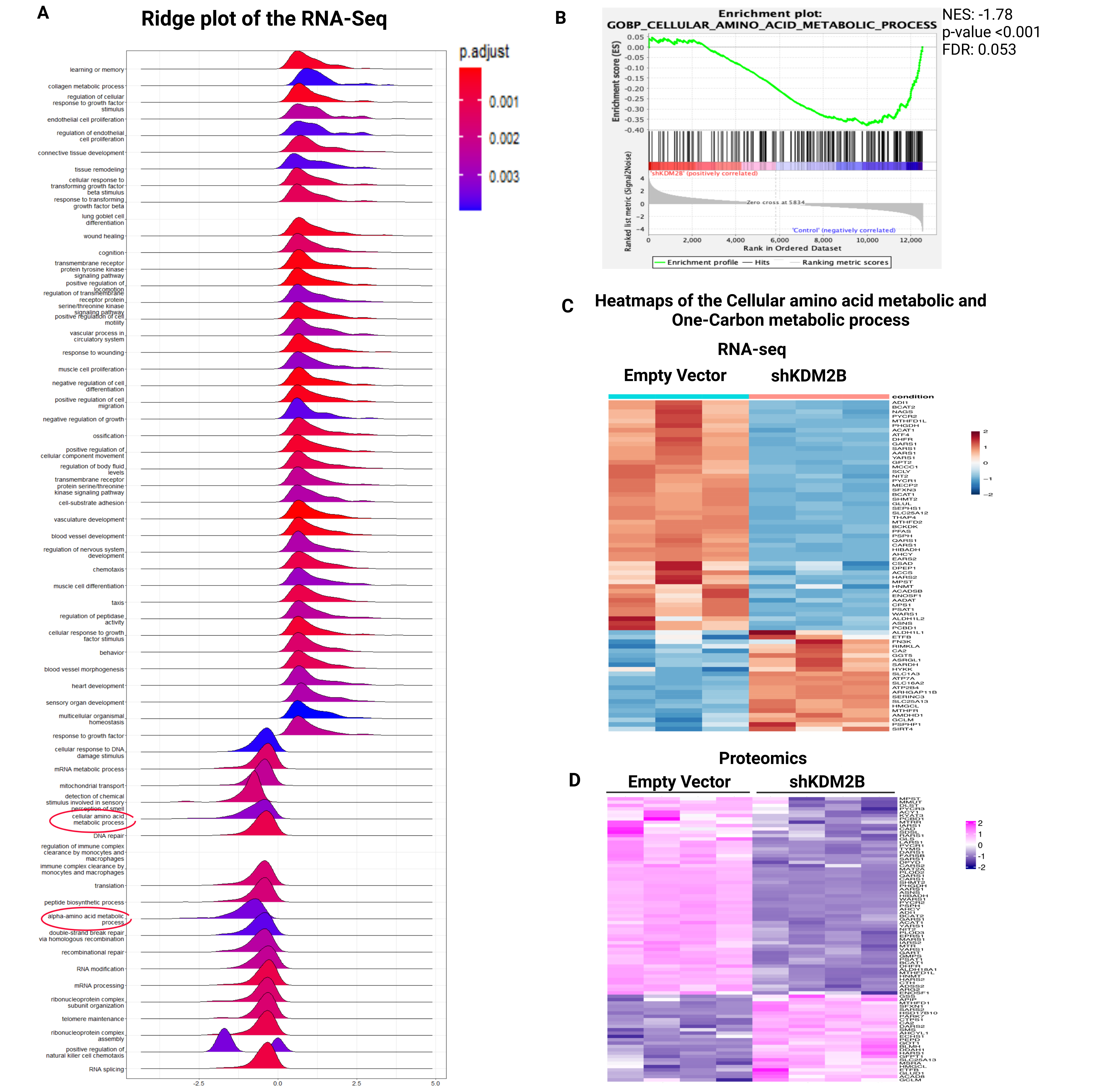

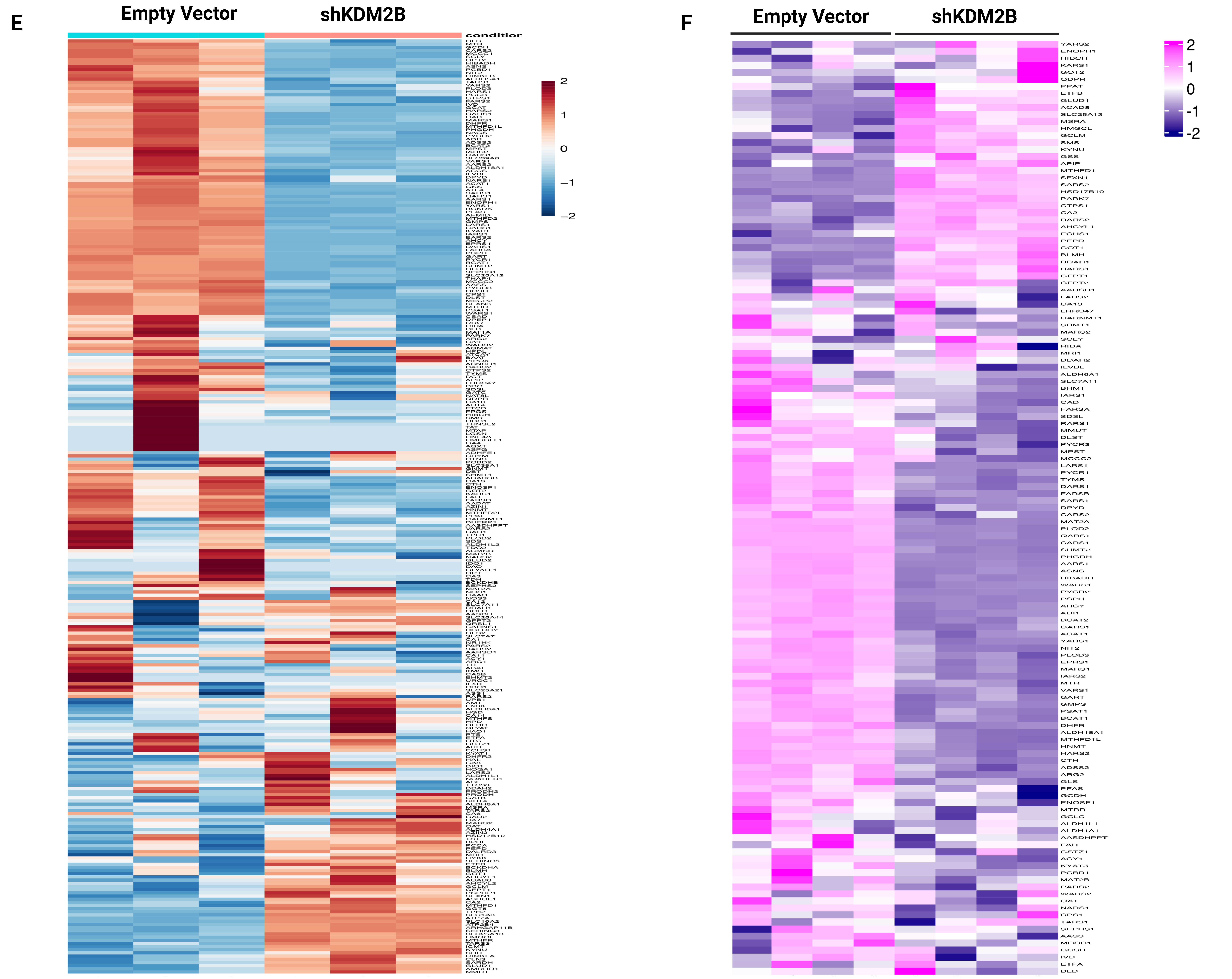


**Supplementary Figure 4. The expression of genes in the GO terms "Cellular amino acid metabolic process" and "One carbon metabolic process" is under the control of KDM2B.**

**A)** Ridge plot analysis of the RNA-seq data, based on all genes in the GO Domain "Biological Process". "Cellular amino acid metabolic process" and the related "Alpha amino acid metabolic processes" were among the significantly downregulated processes in shKDM2B cells. **B)** Gene set enrichment analysis (GSEA) of the RNA-seq data also identified "Cellular amino acid metabolic process" among the statistically significant downregulated biological processes in the shKDM2B cells (NES=-1.78, p-value <0.001). **C)** Heatmap of the statistically significant differentially expressed genes in the GO terms "Cellular amino acid metabolic process" and "One-carbon metabolic process" in Empty vector and shKDM2B-transduced MDA-MB-231 cells. **D)** Heatmap showing the abundance of proteins encoded by genes in the GO terms "Cellular amino acid metabolic process" and "One-carbon metabolic process" in Empty vector and shKDM2B-transduced MDA-MB-231 cells. Protein abundance was measured with quantitative TMT-proteomics. The heatmap shows only proteins whose abundance differs significantly between Empty Vector and shKDM2B cells. **E)** Heatmap of all the differentially expressed genes in the GO terms "Cellular amino acid metabolic process" and "One-carbon metabolic process" in Empty vector and shKDM2B-transduced MDA-MB-231 cells. **F)** Heatmap showing the abundance of proteins encoded by genes in the GO terms "Cellular amino acid metabolic process" and "One-carbon metabolic process" in Empty vector and shKDM2B-transduced MDA-MB-231 cells. The heatmap shows all the proteins whose abundance differs between Empty Vector and shKDM2B cells, including the ones whose difference in abundance is not statistically significant.

#### Supplementary Figure 5.

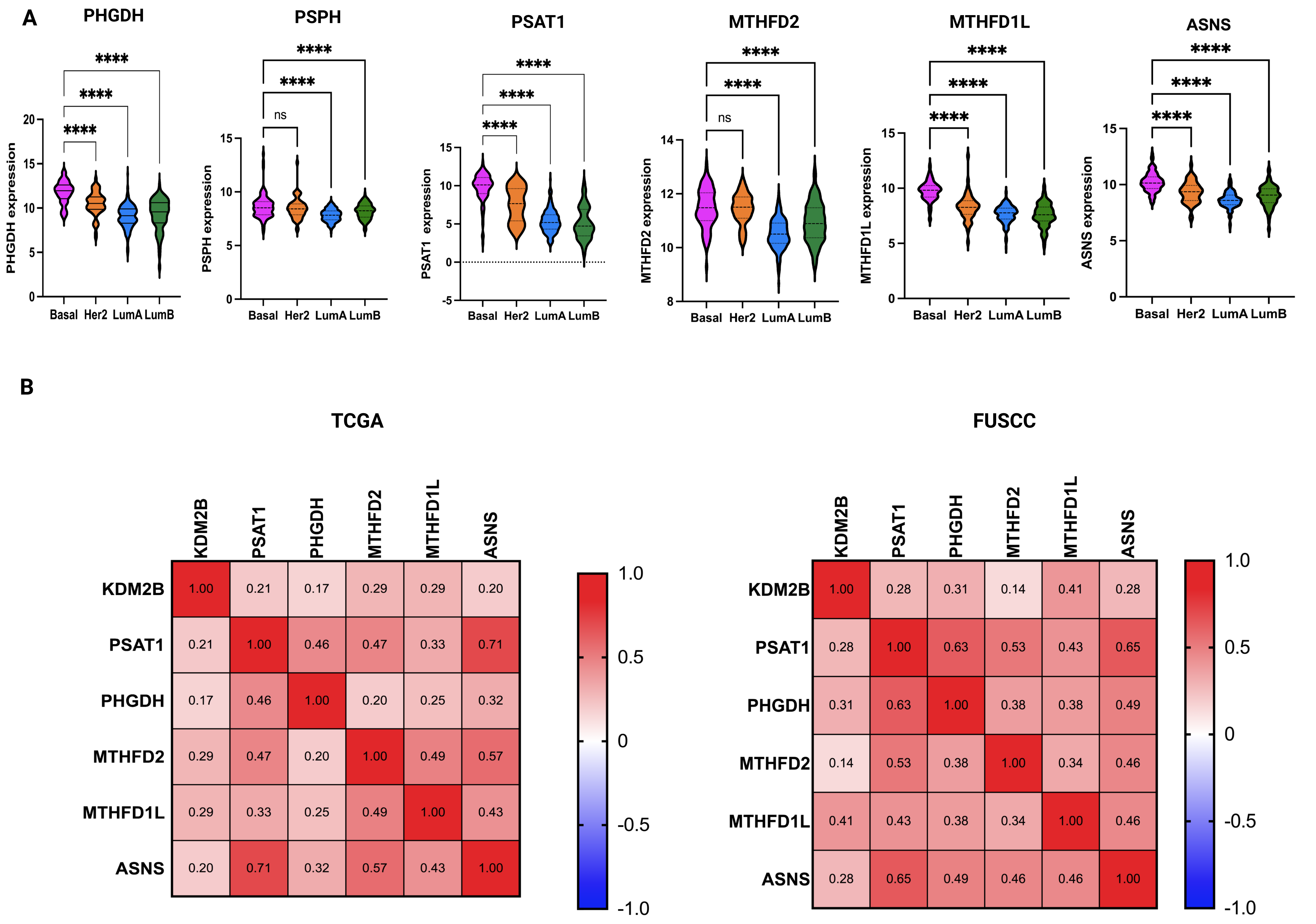


**Supplementary Figure 5. The expression of a set of genes encoding rate limiting enzymes in the SGOC pathway is highest in basal-like breast cancer and correlates with the expression of KDM2B.**
**A)** The RNA levels of the SGOC genes: *PHGDH, PSPH, PSAT1, SHMT2, MTHFD2* and *MTHFD1L* are significantly higher in basal-like TNBC than in Her2, Luminal A and Luminal B breast cancer. Data downloaded from cBioPortal. Statistical significance was determined by One-way ANOVA.**B)** Matrix showing the Spearman correlation of KDM2B expression with the expression of genes encoding rate limiting enzymes in the SGOC or glutamate pathway in 116 TNBC (TCGA data) (left panel) or in 360 TNBC patients (FUSCC cohort) (right panel).

#### Supplementary Figure 6.

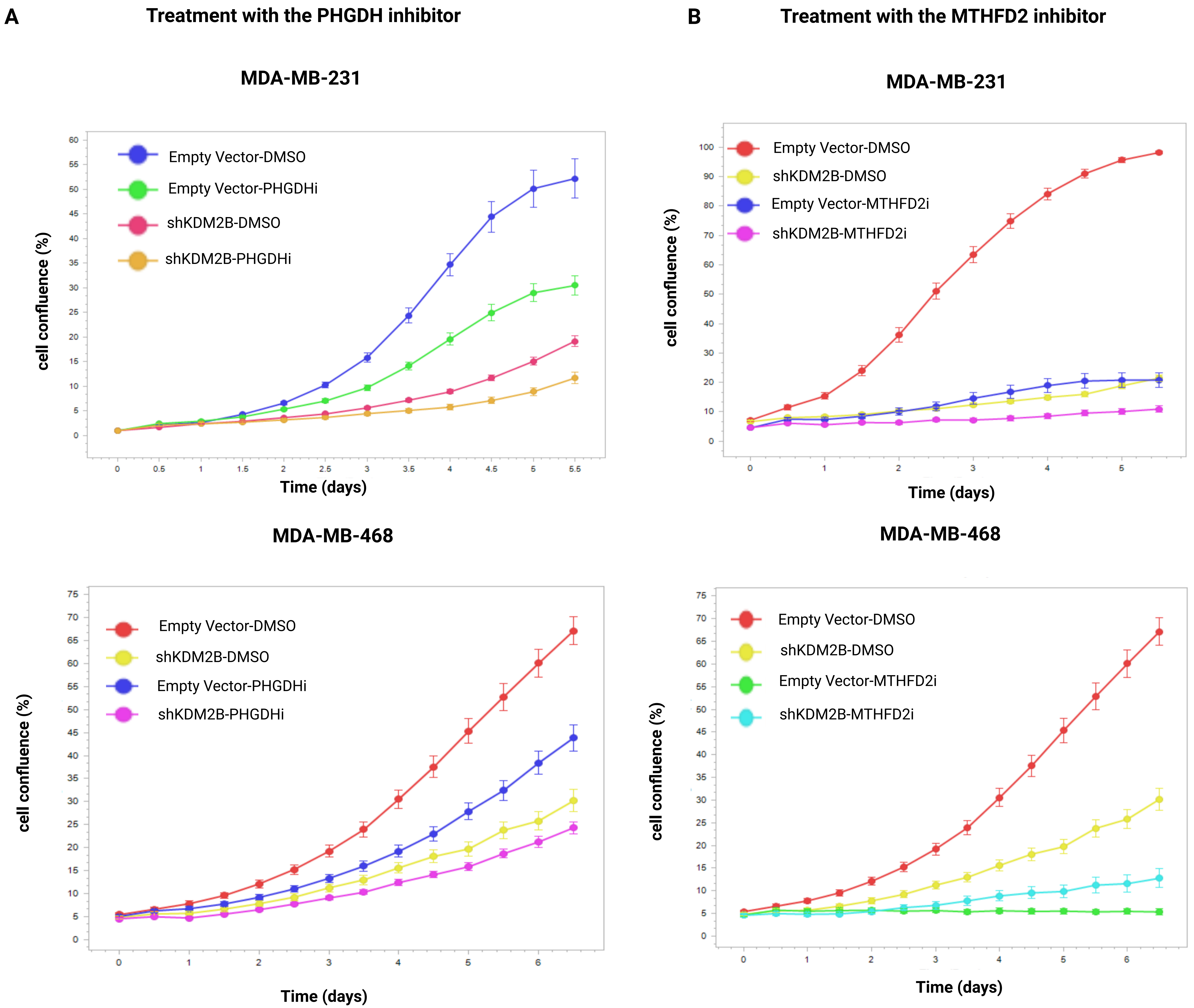


**Supplementary Figure 6. The combination of the knockdown of KDM2B with the pharmacological inhibition of PHGDH or MTHFD2 arrests almost completely the proliferation of both MDA-MB-231 and MDA-MB-468 cells.**
**A)** Growth curves of MDA-MB-231 (top panel) and MDA-MB-468 (bottom panel) cells transduced with the Empty Vector or shKDM2B and treated with 10 μM of the PHGDH inhibitor, NCT-503. Cell confluence was monitored for 5/6 days with the Incucyte live-cell imaging and analysis system. The data shown are from triplicate cultures (n=3). **B)** Growth curves of MDA-MB-231 (top panel) and MDA-MB-468 (bottom panel) cells transduced with the Empty Vector or shKDM2B and treated with 10 μM of the MTHFD2 inhibitor, (DS18561882). Cell confluence was monitored for 5/6 days with the Incucyte live-cell imaging and analysis system. The data shown are from triplicate cultures (n=3).

#### Supplementary Figure 7.

A

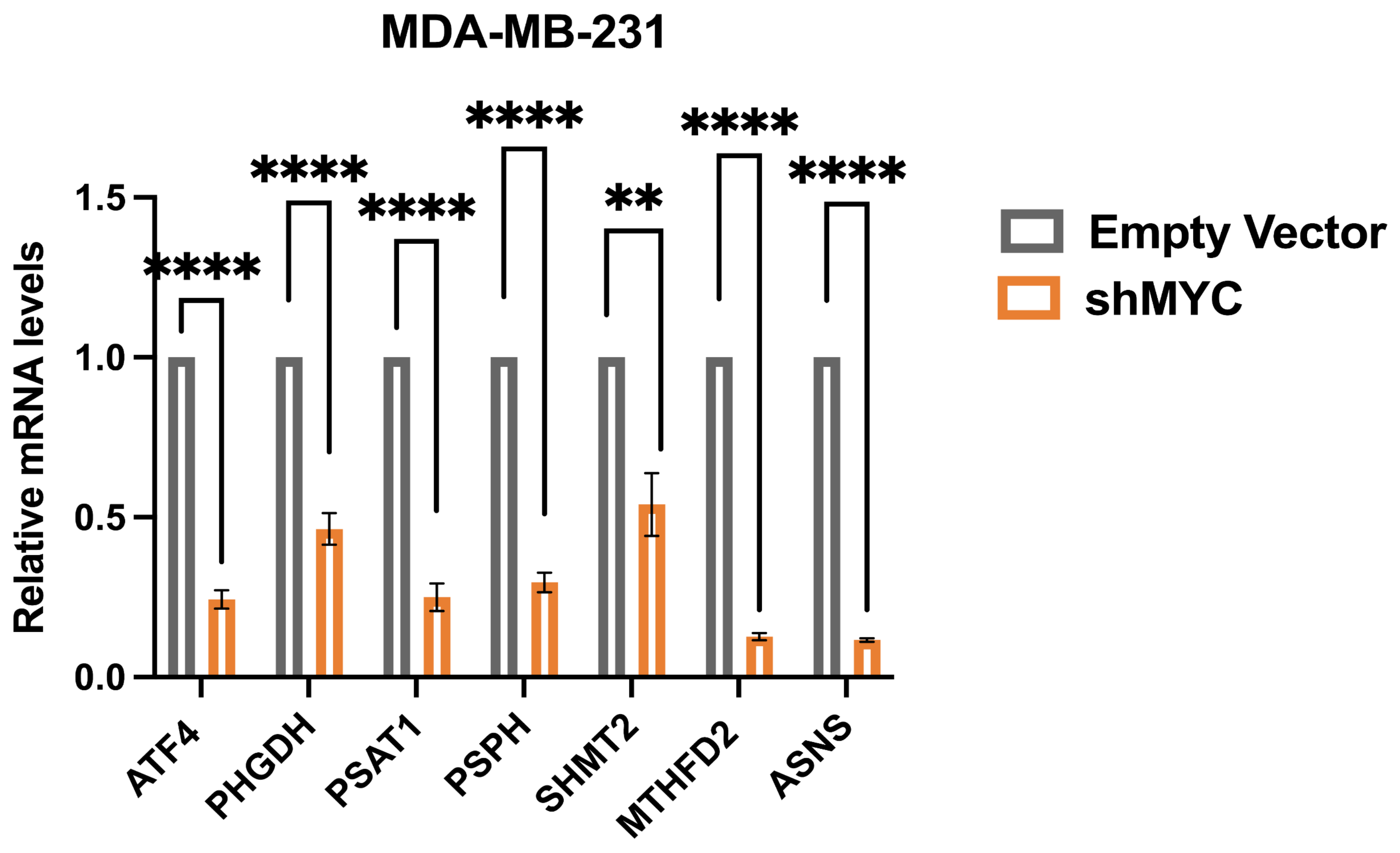


B

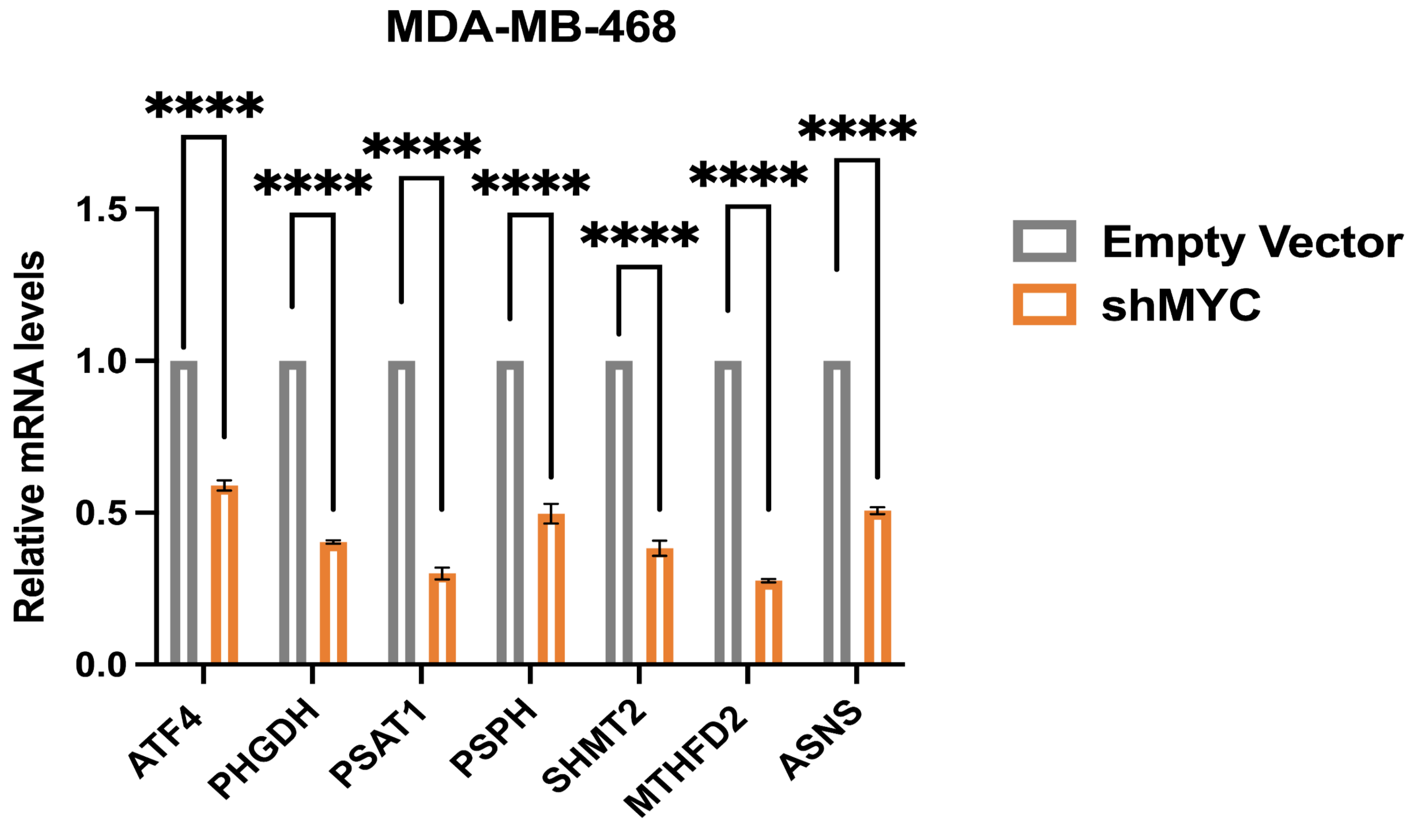


**Supplementary Figure 7. Epistatic relationship of KDM2B, MYC and ATF4.**

**A)** Relative m-RNA levels of ATF4, PHGDH, PSAT1, PSPH, SHMT2, MTHFD2 and ASNS in Empty Vector and shMYC-transduced MDA-MB-231 cells, measured by qRT-PCR. The loading control was β-actin. Mean ± SD of three replicates (n=3). Statistical significance was determined with the unpaired two-tailed t-test. **B)** Relative m-RNA levels of ATF4, PHGDH, PSAT1, PSPH, SHMT2, MTHFD2 and ASNS in Empty Vector and shMYC-transduced MDA-MB-468 cells, measured by qRT-PCR. The loading control was β-actin. Mean ± SD of three replicates (n=3). Statistical significance was determined with the unpaired two-tailed t-test.

**Supplementary Figure 8.**

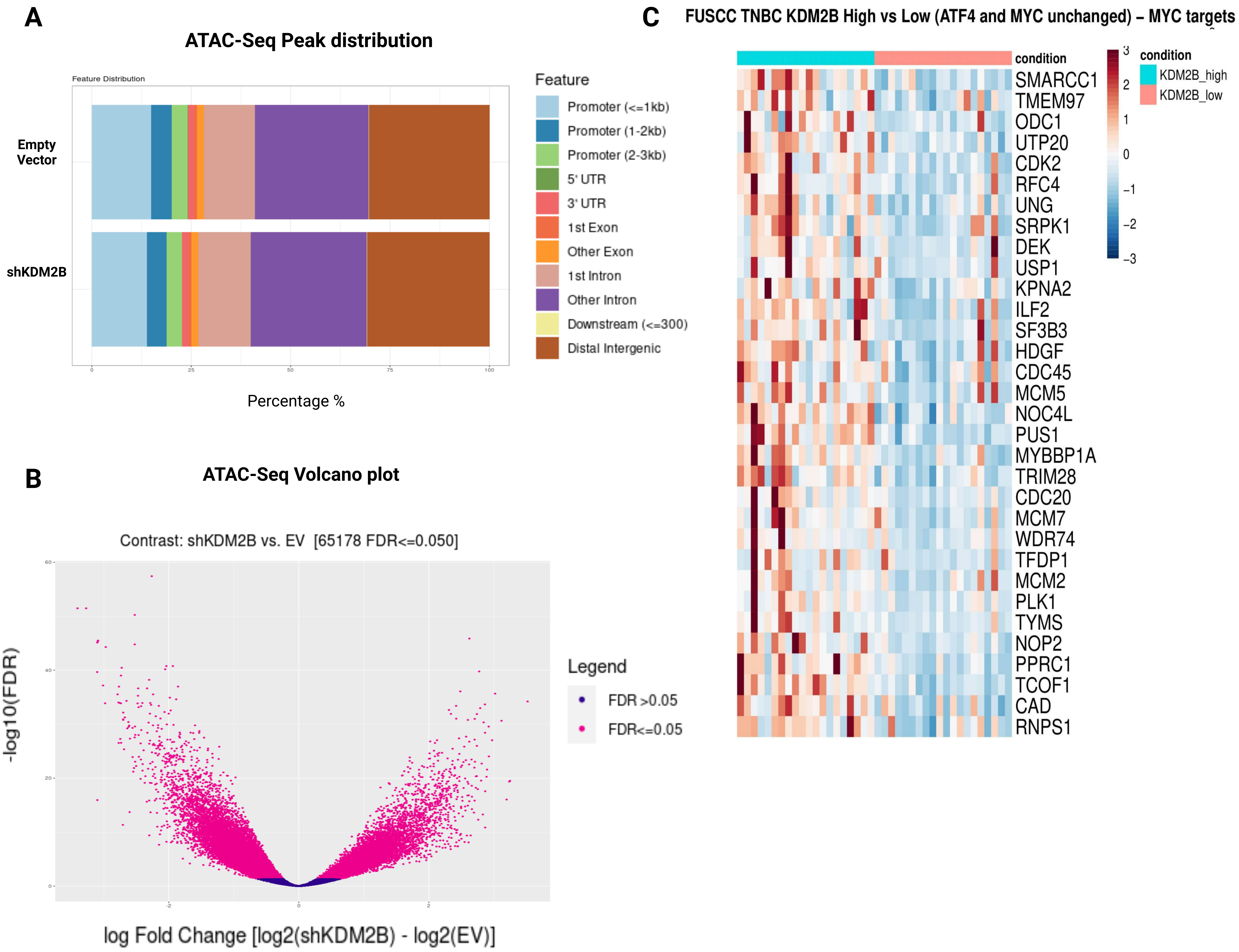


**Supplementary Figure 8. Effect of KDM2B on chromatin accessibility.**
**A)** Most peaks are in the region that extends up to 3 kb upstream of the TSS and includes the first exon and the first intron. Large numbers of peaks were also detected in distal intergenic regions, and in introns other than the first. The knockdown of KDM2B results in a slight reduction of peaks located up to 1 kb upstream from the TSS, but it does not significantly affect peak distribution.**B)** Volcano plot showing the shKDM2B-induced changes in chromatin accessibility in MDA-MB-231 cells. Chromatin accessibility genome-wide was measured by ATAC-seq, which was performed on three biological replicates. The x axis shows the log2 fold change and the y axis the −log10 (FDR). Genes with significant changes in chromatin accessibility (FDR<=0.05) are represented by magenta dots. **C)** Heatmap of the expression of MYC target genes (Hallmarks of V1 and V2 targets, including 240 genes) in a subset of the FUSCC TNBC tumors expressing high or low levels of KDM2B but similar levels of MYC and ATF4. The expression of a subset of MYC target genes (32/240, 13.3% of the genes) was reduced in TNBCs expressing low levels of KDM2B.

### Supplementary Figure 9.

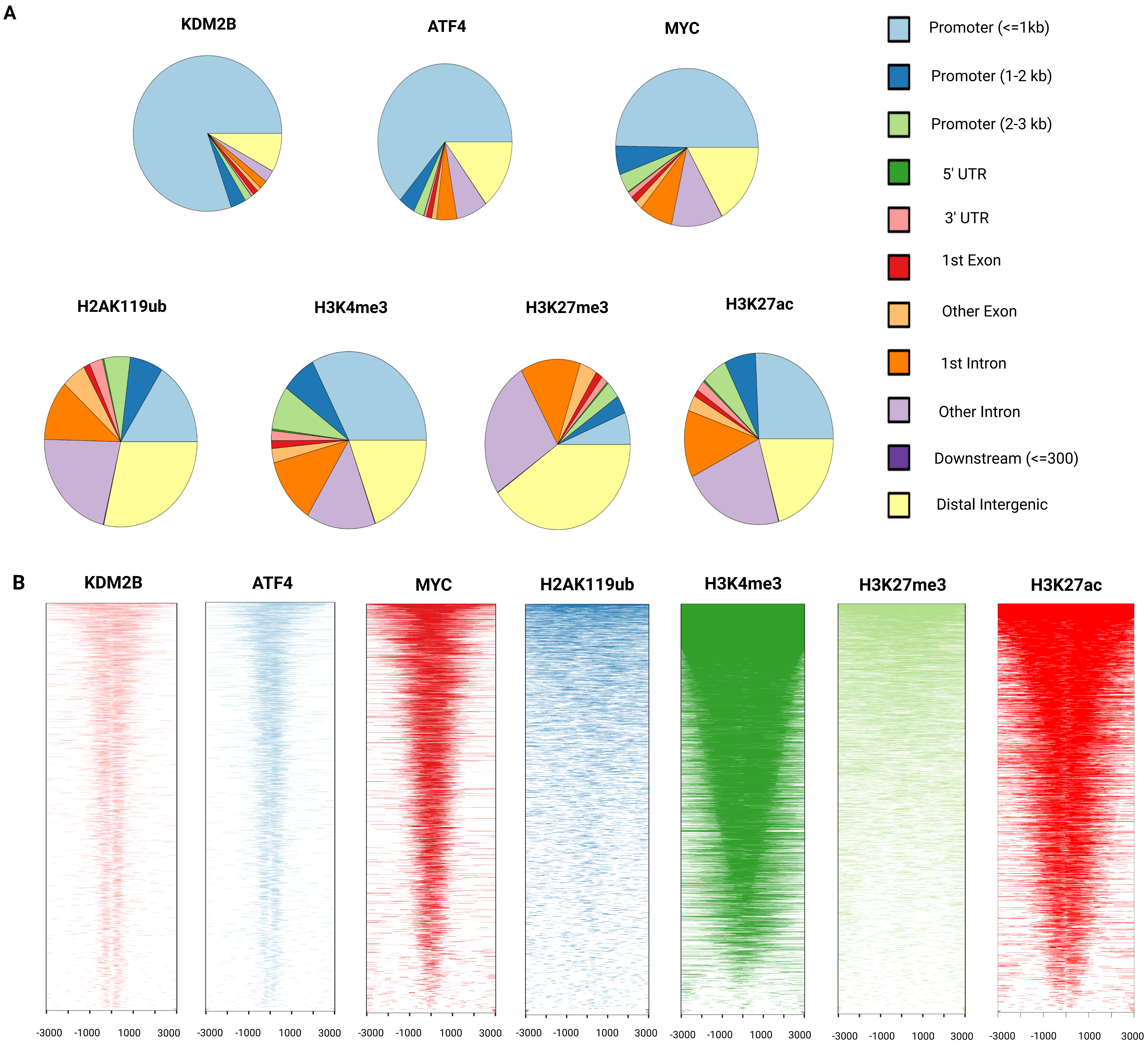


**Supplementary Figure 9. Co-regulation of transcriptionally active genes by KDM2B MYC and ATF4.**
**A)** Genome-wide distribution of ChIP-Seq-detected KDM2B, ATF4, MYC, H2AK119Ub, H3K4me3, H3K27me3 and H3K27ac peaks in MDA-MB-231 cells.
**B)** Heatmaps showing the distribution of ChIP-Seq-detected peaks of KDM2B, ATF4, MYC, and the histone marks listed above, around the TSS.
