## Supplementary Tables of the Methods for "Transcriptional regulation of amino acid metabolism by KDM2B, in the context of ncPRC1.1 and in concert with MYC and ATF4"

**Supplementary Table S10. Lentiviral shRNAs**

| **Gene** | **TRCN clone ID** | **Company** |
| --- | --- | --- |
| KDM2B (#37) | TRCN0000118437 | Millipore-Sigma |
| KDM2B (#79) | TRCN0000238779 | Millipore-Sigma |
| ATF4 | TRCN0000013573 | Millipore-Sigma |
| c-MYC | TRCN0000039640 | Millipore-Sigma |
| EZH2 | TRCN0000040075 | Millipore-Sigma |
| BMI1 | TRCN0000020157 | Millipore-Sigma |
| PCGF1 | TRCN0000073128 | Millipore-Sigma |
| RING1B | TRCN0000033695 | Millipore-Sigma |
| BCOR | TRCN0000033461 | Millipore-Sigma |
| pLKO.1 Empty Vector (TRC Control) | 10879 | Addgene |

**Supplementary Table S11. Treatments**

| **Type of treatment (inhibitors, ROS inducer, scavengers)** | **Cat Number** | **Company** |
| --- | --- | --- |
| Piperlongumine (PL) | HY-N2329 | MedChemExpress |
| APR-246 (Eprenetapopt) | [HY-19980](https://www.medchemexpress.com/PRIMA-1Met.html) | MedChemExpress |
| MI-2 (MALT1 inhibitor) | HY-12276 | MedChemExpress |
| N-Acetyl-L-cysteine (NAC) | A9165 | Millipore-Sigma |
| Glutathione ethyl ester (GSHee) | 14953 | Cayman |
| PHGDH inhibitor (NCT-503) | HY-101966 | MedChem Express |
| MTHFD2 inhibitor | [HY-130251](https://www.medchemexpress.com/ds18561882.html) | MedChem Express |
| Tunicamycin | T7765 | Millipore-Sigma |

**Supplementary Table S12. qRT-PCR primers**

| **Gene** | **Forward** | **Reverse** |
| --- | --- | --- |
| C-MYC | TGGTGCTCCATGAGGAGA | CCAGCAGAAGGTGATCCAGAC |
| PHGDH | GCAAAGAGGAGCTGATAGCG | TTCTCAGCTGCGTTGATGAC |
| PSAT1 | TGCCGCACTCAGTGTTGTTAG | GCAATTCCCGCACAAGATTCT |
| PSPH | GAGGACGCGGTGTCAGAAAT | GGTTGCTCTGCTATGAGTCTCT |
| SHMT2 | CCCTTCTGCAACCTCACGAC | TGAGCTTATAGGGCATAGACTCG |
| MTHFD2 | CTGCGACTTCTCTAATGTCTGC | CTCGCCAACCAGGATCACA |
| ASNS | CAGAAGATGGATTTTTGGCTG | TGTCCAGGAAGAAAAGGCTC |
| ATF4 | GTCCCTCCAACAACAGCAAG | CTATACCCAACAGGGCATCC |
| B-ACTIN | CAACCGCGAGAAGATGACC | ATCACGATGCCAGTGGTACG |

**Supplementary Table S13. Antibodies**

| **Antibody** | **Cat Number** | **Company** | **Application** |
| --- | --- | --- | --- |
| KDM2B | 09-864 | Millipore | WB |
| KDM2B | 17-10264 | Millipore-Sigma | ChIP |
| Ubiquitin (P37) | 58395 | Cell Signaling Technology | WB |
| PHGDH | HPA021241 | Millipore-Sigma | WB |
| PSPH | HPA020376 | Millipore-Sigma | WB |
| PSAT1 | ab154055 | Abcam | WB |
| CTH | 19689 | Cell Signaling Technology | WB |
| SHMT2 | 33443 | Cell Signaling Technology | WB |
| MTHFD1L | 14998 | Cell Signaling Technology | WB |
| MTHFD2 | 41377 | Cell Signaling Technology | WB |
| ALDH1L2 | 21391-1-AP | Proteintech | WB |
| ASNS | 14681-1-AP | Proteintech | WB |
| BCAT1 | 12822 | Cell Signaling Technology | WB |
| BCAT2 | 79764 | Cell Signaling Technology | WB |
| c-MYC | 18583 | Cell Signaling Technology | WB and ChIP |
| ATF4 | 11815 | Cell Signaling Technology | WB and ChIP |
| Phospho mTOR (Ser2481) | 2974 | Cell Signaling Technology | WB |
| mTOR | 2972 | Cell Signaling Technology | WB |
| Phospho eIF2a (Ser51) | 9721 | Cell Signaling Technology | WB |
| eIF2a | 9722 | Cell Signaling Technology | WB |
| EZH2 | 5246 | Cell Signaling Technology | WB |
| BMI1 | 6964 | Cell Signaling Technology | WB |
| PCGF1 | PA549390 | Thermo Scientific | WB |
| RING1B | 5694 | Cell Signaling Technology | WB |
| BCOR | 63972 | Cell Signaling Technology | WB |
| USP7 (HAUSP) | 4833 | Cell Signaling Technology | WB |
| RYBP | 41787 | Cell Signaling Technology | WB |
| TRIM27 | 15099 | Cell Signaling Technology | WB |
| SKP1 | 2156 | Cell Signaling Technology | WB |
| p-H2AX | 9718 | Cell Signaling Technology | WB |
| H2AX | 7631 | Cell Signaling Technology | WB |
| Cleaved PARP | 5625 | Cell Signaling Technology | WB |
| GCLC | ab53179 | Abcam | WB |
| GCLM | 14241-1-AP | Proteintech | WB |
| GSS | 15712-1-AP | Proteintech | WB |
| GSR | ab137513 | Abcam | WB |
| GPX4 | 59735 | Cell Signaling Technology | WB |
| SLC7A11 | 12691 | Cell Signaling Technology | WB |
| GLS1 | 56750 | Cell Signaling Technology | WB |
| GLS2 | 85934 | Cell Signaling Technology | WB |
| β-ACTIN | 4970 | Cell Signaling Technology | WB |
| H3 | 4499 | Cell Signaling Technology | WB |
| H3 | 4620 | Cell Signaling Technology | ChIP |
| A-actinin | 6487 | Cell Signaling Technology | ChIP |
| H3K4me3 | 9751 | Cell Signaling Technology | ChIP |
| H3K27me3 | 9733 | Cell Signaling Technology | ChIP |
| H2AK119ub | 8240 | Cell Signaling Technology | ChIP |
| H3K27ac | 8173 | Cell Signaling Technology | ChIP |

**Supplementary Table S14. Assays**

| **Type of assay** | **Cat Number** | **Company** | **Notes** |
| --- | --- | --- | --- |
| CellROX | C10493 | Thermo Scientific | ROS levels using FACS |
| Glutathione (GSH) | ab205811 | Abcam | Fluorometric Assay using plate reader |
| Serine | ab241027 | Abcam | Fluorometric Assay using plate reader |
| Glycine | ab211100 | Abcam | Fluorometric Assay using plate reader |
| Formate | MAK059 | Millipore-Sigma | Colorimetric Assay  using plate reader |
| NADPH | ab186031 | Abcam | Colorimetric Assay  using plate reader |
| Glutamate | ab83389 | Abcam | Fluorometric Assay Kit  using plate reader |
| S-Adenosylmethionine (SAM) | MET-5152 | Cell Biolabs | ELISA kit using plate reader |
| XF Cell Mito Stress Test Kit | 103015-100 | Agilent Technologies | XF24 Seahorse |
| Oxyblot Protein Oxidation Detection Kit | S7150 | Millipore-Sigma | Oxidised proteins using WB |
| Annexin-V | ab14085 | Abcam | Apoptosis using FACS |

**Supplementary Table S15. ChIP-qPCR primers**

| **Gene** | **Forward** | **Reverse** |
| --- | --- | --- |
| ATF4 -1075 | GGAGACGGCAGCGAAGAA | TCATCCGGCTCAAGTGCAA |
| ATF4 -637 | CTGGTCCCTGAGGCCACTAA | GGTCTCGGGTCGCTGCTA |
| ATF4 -75 | GGCGGAAGGATGCGTCTGT | GGTGGCCGTGGACCCTGA |
| MYC -204 | CTCTGGAACAGGCAGACACA | GACTGAGTCCCCCAATTTGC |
| MYC -51 | AGCTGTGCATACATAATGCA | GAAAACGATGCCTAGAATG |
| MYC +217 | GGCGCCCTGCAGCCTGGTA | GTTTGACAAACCGCATCCTTG |
